# GraphHDBSCAN*: Graph-based Hierarchical Clustering on High-dimensional Spaces with Applications to Single-cell RNA Sequencing Data

**DOI:** 10.64898/2026.03.24.713924

**Authors:** Seyed Ardalan Ghoreishi, Aleksandra Weronika Szmigiel, James Nagai, Ivan G. Costa, Arthur Zimek, Ricardo J. G. B. Campello

## Abstract

Single-cell RNA sequencing (scRNA-seq) is widely used to resolve cellular heterogeneity across thousands to millions of cells. A major challenge is identifying biologically meaningful cell populations while preserving their hierarchical organization, because broad cell types often split into more specialized subtypes. However, state-of-the-art approaches mostly focus on flat partitions and ignore the hierarchical nature of single-cell data. Here we introduce GraphHDBSCAN*, a graph-based extension of HDBSCAN* with an effectively hyperparameter-free hierarchical clustering stage that operates on a graph representation of the data, enabling robust recovery of both hierarchical and non-hierarchical (flat) relationships in very high-dimensional, noisy datasets. We evaluate GraphHDBSCAN* across multiple scRNA-seq datasets and show that it recovers biologically meaningful hierarchies that agree more closely with curated biological references than alternative hierarchical clustering methods, while revealing fine-grained structure, including monocyte subpopulations. In addition, the method yields high-quality flat partitions that outperform widely used community-detection methods.

## Introduction

Single-cell RNA sequencing (scRNA-seq) has emerged as a groundbreaking technology in modern biology, enabling researchers to study gene expression at single-cell resolution. Recent improvements allow measurements of millions of cells in a single experiment, providing an unprecedented view of cellular heterogeneity and tissue complexity^1^. The multifaceted complexity of these datasets creates major challenges, especially for clustering, which aims to group cells with similar expression patterns and reveal biologically meaningful types, states, and lineages. This is far from trivial because scRNA-seq datasets are very high-dimensional and noisy^2^. In addition, many biological processes show natural hierarchical relationships among cell states; that is, broad cell types can be further divided into more specific cell subtypes^3^. As the size of these datasets grows, often beyond one million cells, computational scalability becomes increasingly limiting^4^. However, much more critical is the fact that the presence of thousands of genes (i.e., dimensions of the data space) gives rise to the various manifestations of the so-called *curse of dimensionality*, which poses fundamental challenges for clustering^5^.

So far, a wide range of clustering algorithms has been proposed to address these challenges. Modularity-based community detection algorithms, most notably Louvain^6^ and its improved successor Leiden^7^, have become standard tools in large-scale single-cell analysis frameworks such as `Seurat`^8^ and `SCANPY`^9^, possibly due to their ability to scale to large single-cell datasets, but mainly because they mitigate the curse of dimensionality to a certain extent by working on a similarity graph representation of the data, instead of the original, high-dimensional feature-vector data space. Some drawbacks of these methods include their stochastic nature^10,11^, the need for a previous dimension reduction in the data with PCA and their sensitivity to hyperparameters, such as the optimization function and especially the resolution hyperparameter, whose selection can be particularly challenging in an unsupervised setting, as recently highlighted by Szmigiel et al.^12^. More noticeably, these methods are generally used to produce a single flat partition at a selected resolution and do not explicitly represent the hierarchical relationships between broad cell populations and their more specialized subtypes.

Other graph-based methods have been proposed that can recover hierarchical community structure. AHSCAN and DHSCAN construct agglomerative and divisive graph hierarchies, respectively, using structural similarity between connected nodes^13,14^. However, their original formulations operate on unweighted graphs, such that structural similarity is determined from neighborhood overlap without incorporating the strength of the original graph connections. Consequently, when applied

to weighted similarity graphs, potentially informative differences in edge weights are discarded. More fundamentally, when a flat partition is required, these approaches select a single level of the hierarchy using a modularity-based criterion. Such a global, horizontal selection imposes a common level of granularity across the hierarchy and cannot independently retain clusters that could be most strongly supported at different depths along different branches. Moreover, these methods organize vertices purely according to structural similarity, which in itself does not intrinsically capture a density profile of clusters and noise across different hierarchical levels, something that can be achieved for low- to moderate-dimensional datasets with conventional feature-vector (not graph-based) hierarchical clustering algorithms, in particular, HDBSCAN*^15,16^.

Paris instead operates directly on weighted graphs and constructs an agglomerative hierarchy using a node-pair-sampling-based distance^17^. It therefore preserves edge-weight information and provides a multiscale dendrogram without requiring a predefined resolution hyperparameter. However, the original Paris formulation does not include an intrinsic procedure for automatically extracting a flat partition from the hierarchy; obtaining such a partition requires an additional post-hoc dendrogram-selection step, such as choosing a cut or a target number of clusters. Furthermore, its hierarchy is defined through successive agglomerative merges based on node-pair-sampling probabilities rather than through density connectivity, and all observations remain assigned within the hierarchy regardless of their level of adherence to clusters (or lack thereof), which means even globally outlying observations will be forcibly assigned to clusters rather than being potentially flagged as noise from a density-connectivity perspective.

More recently, scMustree^18^ and Bonsai^19^ have been proposed for hierarchical exploration of single-cell and high-dimensional data. scMustree constructs terminal cell populations through iterative PCA^20^, *k*-nearest-neighbor (kNN) graph construction, and community detection, and subsequently organizes these populations through bottom-up merging based on cluster-specific functional rankings. Although shown to scale to large numbers of cells, its computational cost depends on the number of terminal populations. Bonsai instead reconstructs a maximum-likelihood tree under a probabilistic model of the high-dimensional observations and their measurement uncertainty. Its tree-space search can be computationally demanding, and the authors provide a faster backbone-based mode for very large datasets. Importantly, neither approach constructs a density-based hierarchy directly from a sparse weighted similarity graph while explicitly identifying low-density observations as uncertain (noise), leaving these properties unaddressed by these recent hierarchy-oriented approaches.

Density-based hierarchical methods such as HDBSCAN*^15,16^ are appealing alternatives to these methods because they can identify clusters of highly varying densities, detect noise observations in a non-stochastic manner, and automatically determine the number of clusters. They are also well suited to recovering clusters with irregular shapes and to representing data at multiple levels of granularity. However, applying HDBSCAN* directly to high-dimensional data is fundamentally challenging because the method relies on reliable non-parametric estimation of local density in the input space. Distances derived from nearest-neighbor relationships provide local density information that is subsequently aggregated to build a density-based clustering hierarchy^15,16^. In high-dimensional feature-vector spaces, however, any finite-size dataset becomes exponentially less representative of the underlying density distribution within that space as dimensionality increases, and nearest-neighbor relationships become less discriminative and more sensitive to noisy or irrelevant dimensions, making non-parametric local density estimation increasingly difficult^5^. Dimensionality reduction such as PCA^20^ can alleviate this problem, but density is then estimated in the projected space, and the resulting hierarchy consequently depends on how the original neighborhood structure is transformed. These limitations help explain the relatively low adoption of such methods in single-cell analysis despite their appealing conceptual properties.

To fill this gap, we introduce GraphHDBSCAN*, a graph-based, density-based hierarchical clustering method designed for very high-dimensional (potentially large-scale, noisy) datasets such as scRNA-seq data. Rather than estimating the density hierarchy directly from pairwise (dis)similarities in the original high-dimensional feature-vector space, GraphHDBSCAN* operates on a sparse similarity graph representation of the data. Such a representation retains locally relevant relationships whereby shared neighborhoods and structural similarities can provide more robust information in high-dimensional settings than primary distances alone^5^. GraphHDBSCAN* exploits this property through the Weighted Structural Similarity (WSS)^21^ transformation, which incorporates both agreement between local neighborhoods and the strengths of their connections before graph-based notions of density and density-connectivity are constructed. The method extends the well-known HDBSCAN* hierarchical clustering method from a feature-vector to a graph-based formulation while preserving its key advantages, including the ability to identify a complete density profile of the data with hierarchically nested clusters and subclusters, recover clusters with arbitrary shapes and highly varying densities across multiple density levels, and flag candidate uncertain observations (noise points in conventional HDBSCAN*). In contrast to widely used graph clustering methods such as Louvain and Leiden, which typically produce a single flat partition for a chosen resolution hyperparameter, GraphHDBSCAN* recovers interpretable density-based hierarchies that can be directly visualized and explored without requiring auxiliary dimensionality reduction techniques such as PCA^20^, t-SNE^22^, or UMAP^23^.

A key ingredient of GraphHDBSCAN* is the use of Core-SG^24^, a recent graph sparsification technique with theoretical guarantees. By adapting Core-SG to the context of sparse similarity graphs and leveraging it to efficiently compute clustering hierarchies across a range of density smoothing (a.k.a. *minPts*) values, GraphHDBSCAN* enables the simultaneous efficient computation of a whole family of hierarchies for visual exploration rather than requiring the user to select and tune a single *minPts* value before clustering. This makes the hierarchical clustering stage effectively hyperparameter-free. Notice that the choices involved in constructing the sparse similarity graph provided as input to the algorithm, such as the neighborhood size and similarity measure, belong instead to the preceding data-representation stage. These representation choices are not specific to GraphHDBSCAN*, but are likewise required when graph-based clustering methods such as Louvain or Leiden are applied to feature-vector data. GraphHDBSCAN* can optionally produce high-quality flat partitions derived from the hierarchical results, with fully automatic determination of the number of clusters, and includes an optional density-aware label propagation (LP) strategy that assigns labels to uncertain points according to the same density structure underlying the main clustering algorithm. To illustrate the features of GraphHDBSCAN*, we consider the challenging task of clustering very high-dimensional, noisy single-cell RNA data. Our results show that GraphHDBSCAN* recapitulates biologically meaningful cell-type ontology hierarchies, reveals fine-grained subpopulations, and provides a practical mechanism for handling cells initially identified as uncertain. Quantitative comparisons with a diverse suite of hierarchical baseline competitors further show that GraphHDB-SCAN* achieves superior (overall highest) or highly competitive agreement with the curated biological reference hierarchies on datasets for which such referential hierarchies are available. Benchmarking against state-of-the-art methods also shows that flat partitions derived from GraphHDBSCAN* achieve competitive, often superior performance compared with Louvain, Leiden, and HDBSCAN* across a representative collection of real-world scRNA-seq datasets containing curated biological reference clusters (flat ground-truth labels).

## Results

### Overview of GraphHDBSCAN*

GraphHDBSCAN* is a new graph- and density-based hierarchical clustering method designed for high-dimensional, possibly large-scale, noisy datasets (Fig. 1a). To address the high-dimensionality of the data, GraphHDBSCAN* builds upon previous work showing that secondary (dis)similarity measures based explicitly or implicitly on similarity graphs, such as those based on the notion of Shared Nearest Neighbors (SNN)^25^, can provide more reliable performance than primary base distance measures in high-dimensional settings^5^. Notably, this also supports the common practice of using community detection algorithms, such as Louvain^6^ and Leiden^7^, operating on a sparse similarity graph representation, when clustering high-dimensional biological data.

**Figure 1.**
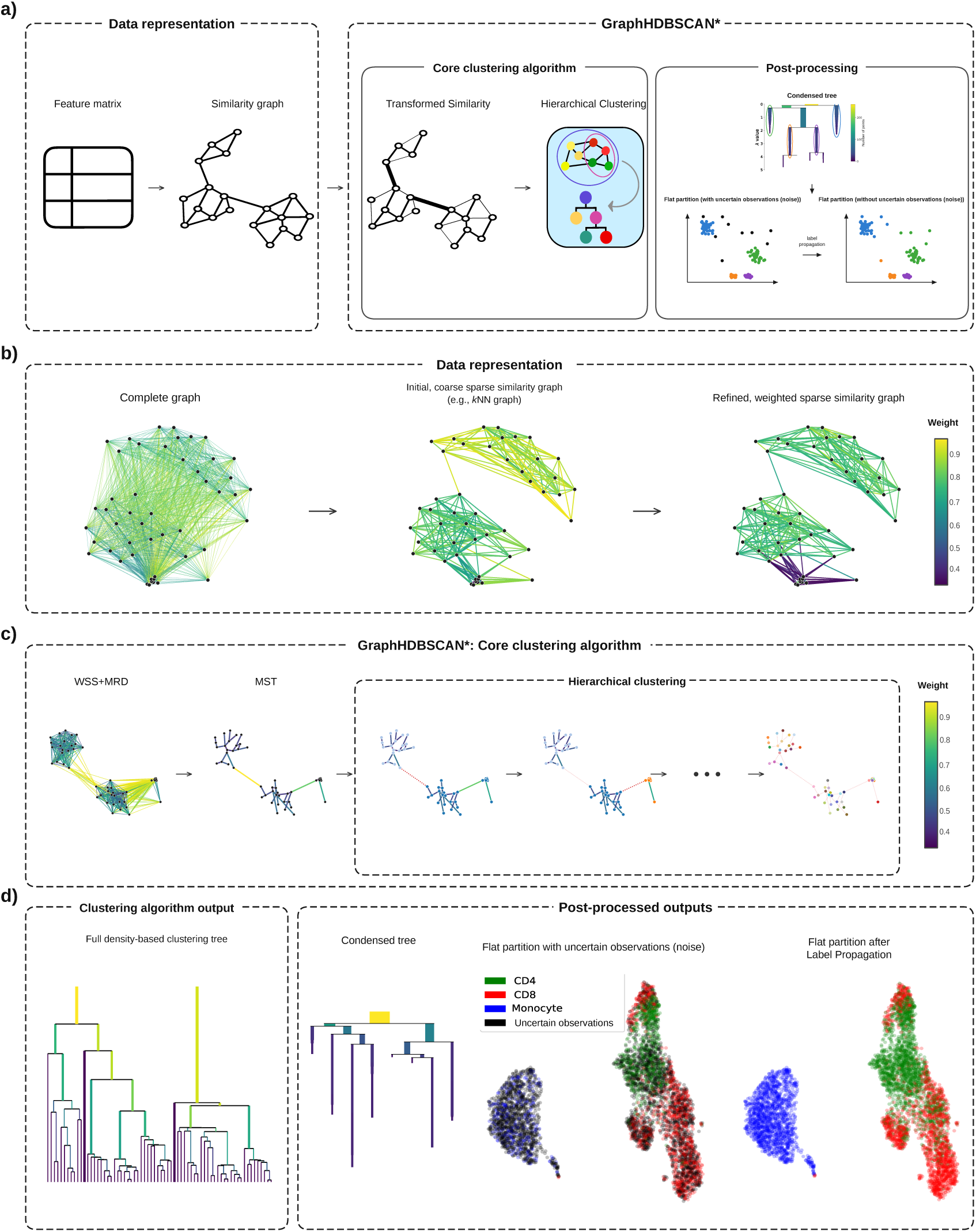
GraphHDBSCAN* workflow. (a) Overview of the complete pipeline. The data representation stage converts the input feature matrix into a sparse weighted similarity graph. The core clustering algorithm transforms the latter graph in order to model density-connectivity and constructs the hierarchical clustering structure, followed by an optional post-processing stage that extracts condensed trees and flat partitions, either retaining explicit uncertain (noise) labels or reassigning them through label propagation. (b) Data representation pipeline. The conceptual complete graph illustrates the underlying pairwise dissimilarities; in practice, an initial, coarse sparse similarity graph (e.g., *k*NN graph) is constructed directly from the feature matrix and then properly reweighted to obtain the final sparse similarity graph. (c) Core GraphHDBSCAN* clustering procedure. The Weighted Structural Similarity (WSS) transformation^21^ is applied to the similarity graph from (b) after which the Mutual Reachability Dissimilarity (MRD) graph is constructed. The corresponding Minimum Spanning Tree (MST) is then obtained and progressively processed to construct the density-based hierarchical clustering structure. (d) Outputs. The core algorithm produces a family of full density-based clustering trees spanning multiple *minPts* values, as the tree in the left panel. Post-processing provides the corresponding condensed trees and for a selected tree, an automatically extracted flat partition that retains observations identified as uncertain. Optionally, label propagation assigns these uncertain observations to clusters.

It is important to emphasize that such a graph-based data representation is an *input* to these algorithms, just as it is an input to GraphHDBSCAN* as well, so its construction should not be confused with (or seen as part of) the clustering algorithms themselves. It should also be noted that there is no unique recipe for constructing a sparse similarity graph. For instance, major single-cell analysis frameworks, including `SCANPY`^9^, `Seurat`^8^, and `PhenoGraph`^26^, follow different variants of a standard pipeline for graph construction, where (Fig. 1b): (a) an initial, coarse sparse similarity graph, such as, e.g., a *k*NN graph (in any of its variant forms), is first constructed directly from feature-vector data — e.g., gene expression data in the form of a feature matrix where rows/observations represent cells and columns/features represent genes, after standard feature selection based on highly variable genes (HVGs); and (b) this initial graph is then properly reweighted and subjected to topological refinements (e.g., to better reflect the weighting scheme and/or to ensure connectedness), giving rise to the final, weighted sparse similarity graph used as input to the clustering algorithms.

Different from other algorithms, which work directly on this input graph, GraphHDBSCAN* derives instead an intermediate, Weighted Structural Similarity (WSS) graph based on a weighted extension of the concept of SNNs^21^, originally defined on binary (unweighted) graphs^25^. It incorporates new edge weights representing similarities between nodes not only directly in terms of explicit adjacency, but also indirectly considering both the topology (shared neighborhoods) and weights of the input graph. The WSS graph allows GraphHDBSCAN* to capture density and density-connectivity in high-dimensional spaces, enabling the algorithm to operate without an additional dimensionality reduction step such as PCA, t-SNE, or UMAP.

Using this graph representation, GraphHDBSCAN* reformulates the building blocks of HDBSCAN*^15,16^, including core distance, mutual reachability, and hierarchy construction, to operate directly on sparse, edge-weighted graphs, thereby producing interpretable density-based cluster hierarchies that would otherwise be hardly detected in high-dimensional spaces. The main output of GraphHDBSCAN* is an entire family of full density-based clustering hierarchies, from which condensed cluster trees can be derived during post-processing. Figure 1 summarizes the complete workflow.

The precursor algorithm, HDBSCAN*, has one mandatory hyperparameter inherited from DBSCAN^27^, namely, *minPts*. It has been shown that HDBSCAN* can be interpreted as being internally performing non-parametric, nearest neighbors-based density estimates, where *minPts* is the number of neighbors and, as such, it controls the smoothness of these estimates as well as that of the resulting hierarchical density profiles^15,16^. Although the algorithm is known to be relatively robust to this hyperparameter, different values of *minPts* may result in different clustering hierarchies^28^. For efficient exploration across different values of *minPts*, we use the Core-SG theoretical machinery to simultaneously derive multiple density-based minimum spanning trees and, consequently, multiple clustering hierarchies for an entire, arbitrary range of *minPts* values. Notably, this can be efficiently achieved from a single initial run with *minPts* = *k*_max_, where *k*_max_ denotes the largest *minPts* value in the desired range. After computing the clustering hierarchy for this single initial run corresponding to the upper-limiting *minPts* value, it can be shown that any other hierarchy corresponding to smaller values of *minPts* can also be extracted nearly instantaneously, with negligible additional cost^24^. In practice, this means that users are not required to select or tune a single *minPts* value before clustering; GraphHDBSCAN* can instead efficiently provide a broad family of density-based hierarchies for direct visualization and exploratory analysis, making the hierarchical clustering stage effectively hyperparameter-free with respect to *minPts*.

As an optional, secondary output, GraphHDBSCAN* can also derive flat partitions from the resulting hierarchies when non-hierarchical clustering is desired. To this end, GraphHDBSCAN* implements the Framework for Optimal Selection of Clusters (FOSC)^29^ in its state-of-the-art, recently extended version, FOSC-X^30^. It allows local, non-horizontal (i.e., multi-level) selection of non-overlapping clusters based on their stability and persistence across the condensed cluster trees, as measured by the well-established Excess-of-Mass (EOM) criterion also adopted by HDBSCAN*^29,30^. Cluster selection can leave outlying observations not assigned to any clusters; these are flagged as uncertain (noise) instead. When a complete data partition is required, the method provides an optional density-aware label propagation procedure based on the semi-supervised classification method HDBSCAN*(cd,−)^31^. As shown in Fig. 1a, this produces a second flat solution in which the initially detected uncertain observations are assigned to the most appropriate clusters. Note that, after the label propagation, the uncertain points identified by GraphHDBSCAN* remain available to the user and can be used for downstream analyses. GraphHDBSCAN* therefore provides three complementary outputs: a family of complete density-based hierarchies and their corresponding condensed cluster trees spanning multiple *minPts* values; for any selected cluster tree, an automatically extracted flat partition retaining explicit uncertain labels; and, optionally, a complete flat partition obtained after label propagation.

### GraphHDBSCAN* Recovers Biologically Meaningful Cellular Hierarchies

The main output of GraphHDBSCAN* is density-based hierarchical clustering. Here, we examine the biological relationships recovered by GraphHDBSCAN* in two blood-derived single-cell datasets using an expert-curated hierarchy of peripheral blood immune cells^32^ shown in Fig. 2a as a biological reference.

**Figure 2.**
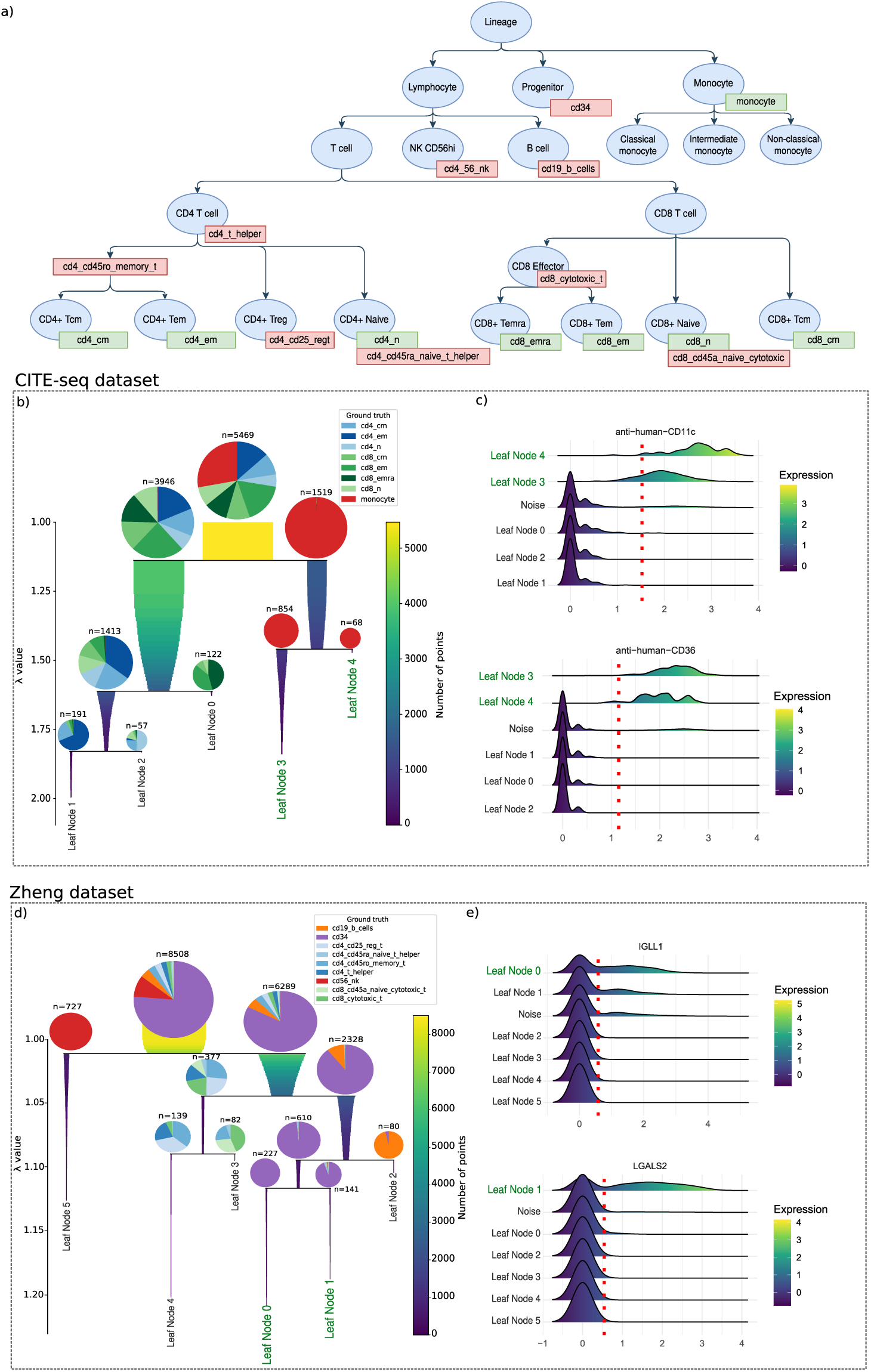
GraphHDBSCAN* accurately reconstructs the curated hierarchical relationships among immune cell types, which can be visualized using a condensed tree. (a) Immune cell lineage tree based on the hierarchy reported by Caron et al.^32^. The green labels indicate the mapping of the cellular annotation described in the CITE-seq annotations, whereas the red labels indicate the mapping of the cellular annotation described in the Zheng annotations. (b) Condensed tree showing the results of GraphHDBSCAN* for the CITE-seq dataset. “Leaf Nodes” represent the clusters associated with the final leaves of the condensed tree. (c) Top markers for the Leaf Node 3 and Leaf Node 4 clusters of the CITE-seq dataset. (d) Condensed tree showing the results of GraphHDBSCAN* for the Zheng dataset. (e) Top markers for the Leaf Node 0 and Leaf Node 1 clusters for the Zheng dataset.

First, we analyzed peripheral blood mononuclear single-cell CITE-seq data from the research conducted by Caron et al.^32^. In CITE-seq, gene expression is simultaneously profiled with protein barcodes, enabling post-sequencing identification of each sequenced cell. Here, we focus only on the RNA-seq modality for clustering and protein antibodies for validation.

As shown in Fig. 2b, the first split of the density-based hierarchy of GraphHDBSCAN* separates monocytes (red) from CD8 (green) and CD4 (blue) T cells. Moreover, the tree can differentiate further between CD8 and CD4 T cells in subsequent splits. Interestingly, when examining the leaves (terminal clusters) in the condensed tree, GraphHDBSCAN* retrieved two monocyte clusters (leaf nodes 3 and 4), which were not described in the reference annotation^32^. Next, we use sc2marker^33^ to identify proteins that can be used to characterize these subgroups with known antibodies. CD36 protein is positive in both leaf nodes 3 and 4, which confirms their monocyte identity^34^. Leaf node 4 shows a higher expression of CD11c, a marker often associated with non-classical monocytes^35^ (Fig. 2c).

Next, we examine the hierarchy in the Zheng dataset^36^. This is a FACS-sorted dataset, so the cell labels come from an orthogonal measurement and can serve as an independent validation of our results. As indicated in Fig. 2d, the first split cleanly separates natural killer cells (red) from all other cell types. Next, we observe a split between CD34+ progenitor and B cells versus CD4+ and CD8+ T cells. These populations are further split at the third level. Here, GraphHDBSCAN* resolves two CD34+ subpopulations, consistent with lymphoid-(IGLL1+) and myeloid-primed (LGALS2+) progenitors, that are not distinguished in the reference labels. The sc2marker analysis indicates that LGALS2 and IGLL1 are cell-surface markers to target when splitting these cell subtypes. While IGLL1+ cells are possibly associated with early pro-B cells^37^, LGALS2+ cells indicate early monocyte cells^38^ (Fig. 2e). Complete marker results, visual comparisons obtained by competing hierarchical clustering approaches, as well as the corresponding analysis of the Bone-marrow^39^ dataset, are provided in the Supplementary Material (Supplementary Figs. S4–S9).

To sum up, GraphHDBSCAN* illustrates the potential of density-based hierarchical clustering as a proxy for recovering meaningful biological signals by identifying the data density backbone, despite its faint signature obscured by the lack of contrast that characterizes high-dimensional, noisy biological data. As an example, by combining GraphHDBSCAN* results with sc2marker, we can select cell-surface markers with validated antibodies to isolate underrepresented cell populations of interest in two blood-derived single-cell datasets. These findings illustrate the potential of GraphHDBSCAN* for dissecting cell populations not described in the original studies producing the data.

### Quantitative Evaluation of Hierarchical Agreement

To complement the qualitative biological analysis presented above, we quantitatively evaluate the agreement between the inferred clustering hierarchies and the corresponding biological reference hierarchies. While standard evaluation measures such as the Adjusted Rand Index (ARI)^40^ and Adjusted Mutual Information (AMI)^41,42^ assess a flat (i.e., non-hierarchical) candidate clustering partition against a flat referential ground-truth partition, here we compute the Hierarchy Agreement Index (HAI)^43^ against the expert-curated cell hierarchy. Higher HAI values indicate greater agreement, with a value of one corresponding to exact agreement between the predicted and reference hierarchies. We use the hierarchy of peripheral blood immune cells^32^ shown in Fig. 2a for the Zheng and CITE-seq datasets. For the Bone-marrow dataset, we use the hematopoietic hierarchy provided in Supplementary Fig. S7. Moreover, these datasets were selected because their cell-type annotations are supported by orthogonal measurements: cell sorting for the Zheng PBMC dataset; protein abundances for the CITE-seq dataset; and cell sorting, protein abundances complemented by atlas based label transfer for the Bone-marrow dataset. We compare GraphHDBSCAN* against the hierarchical clustering methods HDBSCAN*+PCA, HDBSCAN*^15^, Paris^17^, AHSCAN^13^, DHSCAN^14^, scMustree^18^, and Bonsai^19^.

Our hierarchy-based evaluation in Table 1 shows that GraphHDBSCAN* achieves the highest HAI on the Zheng and Bone-marrow datasets, with values of 0.9066 and 0.8331, respectively. On CITE-seq, scMustree^18^ achieves the highest HAI (0.8852), with GraphHDBSCAN* attaining a very similar score of 0.8823. Overall, GraphHDBSCAN* shows consistently strong agreement with the curated biological hierarchies, ranking first on two of the three evaluated datasets and remaining highly competitive on the third. For Bonsai, the available implementation did not complete within one week of runtime on the Zheng and Bone-marrow datasets; these cases are therefore reported as “–”. In contrast, GraphHDBSCAN* took an average of 41.93 seconds in five distinct runs (including the data representation phase) on the largest evaluated dataset (Bone-marrow), with 71,260 cells.

**Table 1.** Agreement between inferred clustering hierarchies and the curated biological reference hierarchies, measured using the Hierarchy Agreement Index (HAI; higher is better). The highest value for each dataset is shown in bold, and the second-highest value in italics. Entries marked “–” indicate cases in which the available implementation did not produce a hierarchy within one week of runtime; these runs were therefore considered computationally prohibitive, and no HAI value could be computed. Further details are provided in the Supplementary Material.

| Method | CITE-seq HAI ↑ | Zheng HAI ↑ | Bone-marrow HAI ↑ |
|---|---|---|---|
| GraphHDBSCAN* | *0.8823* | **0.9066** | **0.8331** |
| HDBSCAN*+PCA | 0.8798 | 0.8654 | 0.7179 |
| HDBSCAN* | 0.8459 | *0.8837* | 0.7256 |
| Paris | 0.8740 | 0.8219 | 0.7518 |
| AHSCAN | 0.8656 | 0.7583 | 0.6366 |
| DHSCAN | 0.7868 | 0.8177 | 0.6638 |
| scMustree | **0.8852** | 0.7462 | *0.7557* |
| Bonsai | 0.8033 | – | – |

Together, these qualitative and quantitative results demonstrate the ability of GraphHDBSCAN* to recover biologically meaningful hierarchical relationships while revealing finer-grained cell subpopulations.

### Benchmarking GraphHDBSCAN* Flat Partitioning Against State-of-the-Art Methods

We next evaluate the flat partitioning performance of GraphHDBSCAN* against the original HDBSCAN*, HDBSCAN* applied after dimensionality reduction using PCA, and a diverse set of established clustering methods, including Louvain, Leiden, Paris^17^, FlowSOM^44^, DHSCAN, AHSCAN, and SC3^45^. Clustering performance is assessed using the ARI and AMI^40–42^ across 17 single-cell datasets from the benchmarking framework proposed in^12^. A key characteristic of these single-cell datasets is that their cell labels were not derived using clustering algorithms, which avoids biasing the evaluation toward methods similar to those used to generate the labels. The benchmark was performed under two controlled evaluation scenarios: (1) Default—using the default SCANPY settings for graph construction where applicable and the default algorithm-specific settings for each clustering method; and (2) Best—using the best-performing configurations. For the Best scenario, the graph-construction settings for graph-based methods and, where applicable, the algorithm-specific hyperparameters were optimized to identify the best-performing configuration for each method. Details on how optimization was performed and how methods were applied are provided in the Methods section.

As illustrated in Fig. 3, GraphHDBSCAN* shows consistently strong performance with respect to both evaluation measures, despite not being primarily designed for flat partition extraction. The boxplots under the default and best-scenario configurations show that GraphHDBSCAN* obtains the highest median ARI and AMI in both evaluation scenarios, with its score distributions remaining concentrated near the upper end of the evaluation range. Under the best-scenario configuration, GraphHDBSCAN* achieves median ARI and AMI scores of 0.9727 and 0.9590, respectively. Among the closest competitors, Louvain achieves median ARI and AMI scores of 0.9587 and 0.9398, respectively, while Leiden reaches 0.9463 and 0.9350. FlowSOM also performs well, with median scores close to those of Louvain and Leiden. The original HDBSCAN* achieves considerably lower median scores of 0.3442 for ARI and 0.4035 for AMI.

**Figure 3.**
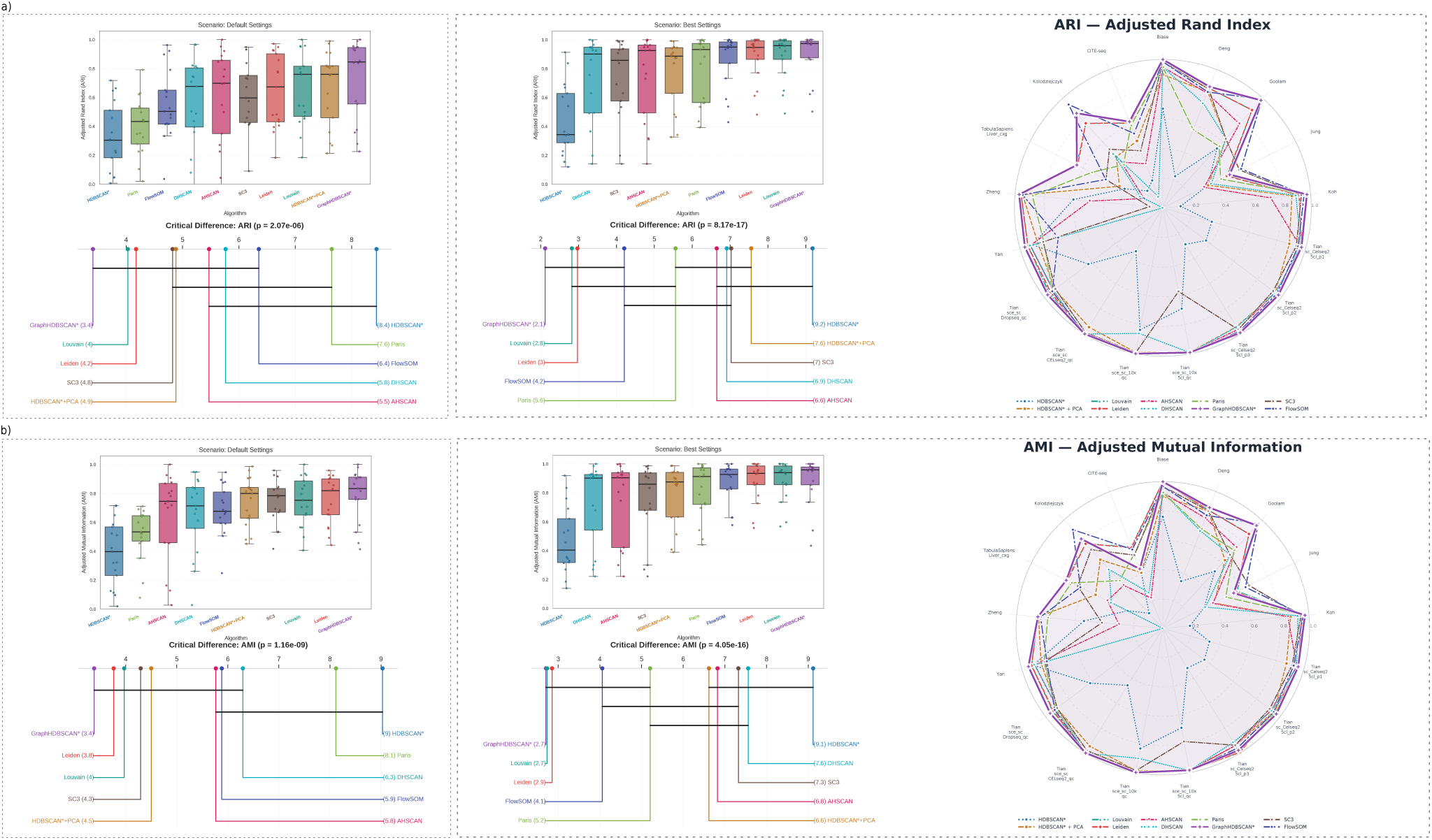
Benchmarking flat partitioning performance across 17 single-cell RNA sequencing datasets. (a) ARI-based evaluation comparing HDBSCAN*, HDBSCAN*+PCA, Louvain, Leiden, GraphHDBSCAN*, Paris, FlowSOM, DHSCAN, AHSCAN, and SC3. Boxplots summarize the distributions of ARI scores across datasets under the default and best-scenario configurations, while the Critical Difference (CD) diagrams report the corresponding average ranks. (b) AMI-based evaluation presented in the same format as in (a), including boxplots and CD diagrams for the default and best-scenario configurations. The radar plots provide a per-dataset comparison of ARI and AMI performance using the best available tuned configuration for each method. The Louvain and Leiden curves largely overlap in the radar plots because their per-dataset performances are identical or very similar across many datasets.

The boxplots in Fig. 3 further indicate that GraphHDBSCAN* performs consistently across datasets rather than deriving its overall performance from only a small number of particularly favorable cases. Notably, the graph-construction stage in the default scenario uses the standard `SCANPY` default settings, without dataset-specific optimization, whereas `SCANPY`, being developed for scRNA-seq analysis, provides Louvain/Leiden defaults that act as a practical baseline for this type of data. Fig. 3 also shows that applying PCA substantially improves HDBSCAN* relative to its direct application to the original high-dimensional data; nevertheless, HDBSCAN*+PCA is still outperformed by the strongest-performing methods under the best-scenario configuration. The original HDBSCAN* exhibits the lowest median performance and the greatest number of low-scoring results, underscoring its limitations when applied directly to high-dimensional single-cell data. Another interesting result is that the mean performance of Leiden, Louvain, FlowSOM, and Paris dropped considerably from the best to the default settings (ARI: 25.2–49.8%; AMI: 14.5–36.9%). In comparison, the drops for GraphHDBSCAN* (ARI: 18.7%; AMI: 11.0%) and HDBSCAN*+PCA (ARI: 15.1%; AMI: 6.7%) were smaller. Notably, GraphHDBSCAN* also achieved the highest mean performance in both scenarios, which highlights both its accuracy and its robustness to hyperparameter choices.

The radar plots provide a per-dataset view of clustering quality under the best-scenario configuration and show that GraphHDBSCAN* maintains strong ARI and AMI performance across the 17 benchmark datasets. For ARI, GraphHDBSCAN* achieves or shares the highest score on 11 of the 17 datasets, including 7 datasets on which it obtains the strictly highest score. For AMI, it achieves or shares the highest score on 8 datasets, including 4 strictly highest scores. GraphHDBSCAN* also attains perfect ARI and AMI values on the Biase, Tian sce_sc_10x qc, and Tian sce_sc CELseq2_qc datasets. Other methods show advantages on particular datasets, reflecting differences in the cluster structure captured by the individual approaches. Nevertheless, GraphHDBSCAN* remains highly competitive across the complete collection of datasets and rarely exhibits substantial drops in performance.

To assess overall performance trends, the CD diagrams summarize the average rankings of the ten methods across all datasets. Statistical differences among methods were assessed using the Friedman test, followed by the Nemenyi post-hoc test for pairwise comparisons^46^. The CD diagrams visualize the average method ranks and the corresponding critical differences obtained from the Nemenyi test. GraphHDBSCAN* obtains the best average rank for default-scenario ARI (3.4), best-scenario ARI (2.1), and default-scenario AMI (3.4), while GraphHDBSCAN* and Louvain share the best average rank of 2.7 for best-scenario

AMI. Louvain and Leiden are among the closest competing methods overall, while FlowSOM also performs particularly well under the best-scenario configurations. The original HDBSCAN* ranks last in every comparison. The corresponding Friedman tests indicate statistically significant overall differences among the evaluated methods for default-scenario ARI (*p* = 2.07 × 10^−6^), best-scenario ARI (*p* = 8.17 × 10^−1^^7^), default-scenario AMI (*p* = 1.16 × 10^−9^), and best-scenario AMI (*p* = 4.05 × 10^−1^^6^). These rankings demonstrate that GraphHDBSCAN* performs consistently at or near the top across datasets, evaluation measures, and evaluation scenarios. The exact numerical values of ARI and AMI are reported in Supplementary Table S2. Moreover, we compare the runtime scaling of GraphHDBSCAN* with Louvain and Leiden across datasets of increasing size. GraphHDBSCAN* shows comparable scaling across the evaluated datasets (Supplementary Fig. S3).

Overall, these results demonstrate that GraphHDBSCAN* provides consistently strong flat partitioning performance across a broad range of single-cell datasets and across a diverse collection of competing clustering approaches. Importantly, this performance is obtained while GraphHDBSCAN* also provides a whole family of hierarchical representations of the data and identifies uncertain observations. The strong performance of GraphHDBSCAN* highlights the effectiveness of incorporating graph structure into HDBSCAN*.

### GraphHDBSCAN* Label Propagation Assigns Labels to Uncertain Points

While a byproduct of GraphHDBSCAN* is the identification of uncertain observations based on data density, real-world applications such as resource-intensive single-cell analyses may benefit from a complete labeling of observations. To provide a complete dataset partitioning, GraphHDBSCAN* employs a label propagation approach. To illustrate the benefits of the GraphHDBSCAN* strategy, we consider the well-known PBMC3k dataset^36^ used in single-cell tutorials.

After applying GraphHDBSCAN*, the flat partition results were annotated using the cell-type markers available in Supplementary Table S3. This enabled the identification of nine cell types, while 1673 of the 2695 cells were classified as uncertain. Uncertain points were subsequently reassigned using the label-propagation approach. As illustrated in Fig. 4a, the majority of points were reassigned to the T cell cluster, with the second-largest proportion assigned to the cDC2 cluster. To further investigate the accuracy of the label-propagated clusters, we identified the top 20 gene markers for each original cluster prior to label propagation. Next, we measured the correlation between the expression profiles of cells originally allocated to the cluster and the label-propagated cells (LP). As shown in Fig. 4b, we observe a higher correlation (0.983±0.012) for matching cell types versus an average correlation of (0.30±0.36) between non-matching cell types.

**Figure 4.**
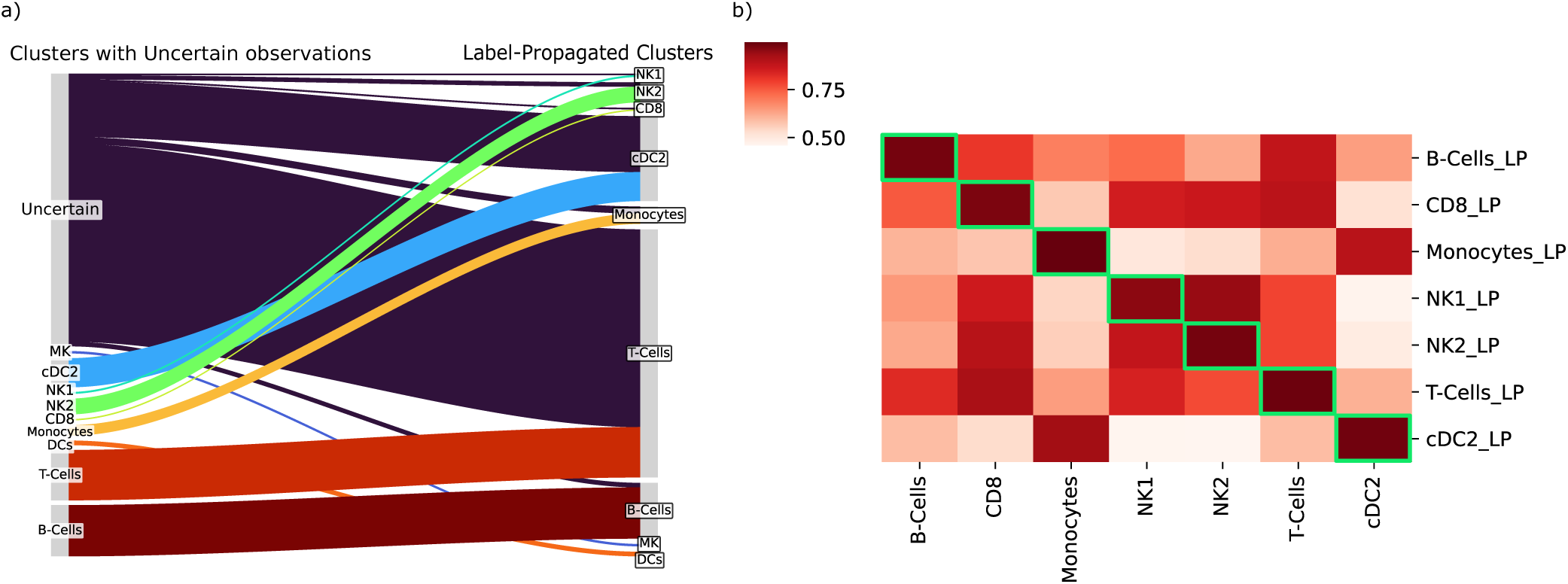
(a) Sankey plot indicating the assignment of clusters before (left) or after (right) label propagation. (b) Pearson correlation matrix of the top 20 features of the clusters (x-axis) and their label-propagated counterparts (y-axis). Green boxes indicate the highest correlation pairs, all of which lie on the diagonal, i.e., for matching cell types.

To further investigate the behavior of label propagation in a dataset with known reference labels, we analyzed the Tian sce_sc_Dropseq_qc dataset^47^. Supplementary Fig. S1 compares the GraphHDBSCAN* assignments before and after label propagation with the reference cell-line labels across PCA, t-SNE, and UMAP representations. The visualization shows that, although PCA, t-SNE, and UMAP do not consistently capture cells from the same reference population within a single group, GraphHDBSCAN* relabels the initially identified uncertain points to their correct labels. Because points identified as uncertain may also include technical artifacts, we additionally examined their relationship with doublet predictions in the same dataset. We observe that the points identified as uncertain by GraphHDBSCAN* (5 out of 225 cells) exhibit higher doublet scores than cells assigned directly to the main clusters (Supplementary Fig. S2). Together, these results illustrate that the GraphHDBSCAN* uncertain points can contain both observations that can be consistently reassigned to reference populations through label propagation and observations associated with potential technical artifacts such as doublets.

## Discussion

A central challenge for density-based clustering in scRNA-seq is high-dimensionality: distances become less informative, which weakens density estimation and makes direct application of HDBSCAN* both less effective and computationally inefficient. In practice, this is often addressed through representation learning or dimensionality reduction (e.g., PCA^20^, t-SNE^22^, UMAP^23^) before clustering. While useful, these steps can reshape the neighborhood and density structure, so clustering results may partly reflect the embedding rather than the original data.

To address these challenges, we propose GraphHDBSCAN*, a clustering method with an effectively hyperparameter-free hierarchical clustering stage that combines graph topology and density directly on a sparse similarity graph, without requiring any auxiliary dimensionality reduction. This HDBSCAN*-style hierarchical clustering stage is both accelerated and made effectively hyperparameter-free by leveraging the Core-SG^24^ machinery, which efficiently derives the family of hierarchies across multiple *minPts* values without requiring the user to select a single *minPts* value before clustering.

In this manuscript, we evaluate GraphHDBSCAN* from two complementary perspectives. First, we provide an evaluation of hierarchical clustering, including seven competing hierarchical clustering methods and three single-cell datasets with expert-curated hierarchies. Second, a flat partitioning benchmark comprising seventeen datasets and nine widely used competing methods is presented.

To enable hierarchy-based clustering evaluation, a known hierarchy is required. While such hierarchies are described in detail in sources like the Cell Ontology, datasets with names standardized according to ontology terms remain, to our knowledge, limited, and the mapping is often not straightforward, particularly for highly complex immune cell types^48^. However, immune cell datasets frequently include additional modalities, such as protein measurements in CITE-seq or cell sorting prior to library preparation, which allow for cell identification independent of gene expression data. Here, we leverage three unique blood cell datasets, which include either protein measurements, physical cell sorting and an accompanying expert-curated cell hierarchy to evaluate the performance of GraphHDBSCAN* and competing hierarchical methods.

The hierarchies recovered by GraphHDBSCAN* agreed closely with curated cell-type annotations, ranking first on two of the three reference hierarchies and within 0.003 HAI of the best method on the third. They thus reflected both broad immune lineages and finer subtype organization. In contrast, dimensionality reduction did not consistently help feature-based density clustering: PCA improved standard HDBSCAN* on CITE-seq but reduced its agreement on Zheng and Bone-marrow. This suggests that building the density hierarchy directly on a sparse similarity graph is a more robust alternative to choosing a dataset-specific embedding.

Because HAI is computed against a curated reference hierarchy, it evaluates agreement with the known cellular organization but does not explicitly reward additional biologically supported substructure that is absent from the reference. The recovered hierarchies also revealed biologically interpretable substructure beyond the original coarse annotations. In the CITE-seq dataset, GraphHDBSCAN* separated monocytes from CD4 and CD8 T cells before resolving the two T cell branches and resolved two monocyte subpopulations not distinguished in the reference annotation. In the Zheng dataset, the hierarchy first separated natural killer cells, subsequently distinguished progenitor and B cell populations from T-cell populations, and identified two previously unreported CD34-associated subpopulations characterized by IGLL1 and LGALS2. These findings illustrate the potential of the density hierarchy to generate biologically meaningful hypotheses and suggest markers for the characterization and isolation of potentially undescribed cell populations.

Although GraphHDBSCAN* is primarily a hierarchical method, it also produced consistently strong flat partitions, matching or outperforming Louvain and Leiden on most datasets according to ARI and AMI and clearly improving over direct HDBSCAN* and HDBSCAN*+PCA overall. Across the clustering benchmark datasets, GraphHDBSCAN* achieved the highest median ARI and AMI under both the default and best-scenario settings. The Critical Difference analyses further support the consistency of these results. Moreover, we observe that the drop in ARI and AMI when contrasting the default and best scenario is lower for GraphHDBSCAN* than for Louvain, Leiden, FlowSOM, and Paris, suggesting lower sensitivity to graph-construction and hyperparameter choices.

Notably, this flat partition is extracted using the EOM criterion within FOSC/FOSC-X, which is the standard flat-cluster extraction criterion used by HDBSCAN*. We deliberately retain this standard criterion in our experiments rather than introducing a graph-specific extraction criterion, allowing the contribution of the graph representation and graph-based clustering strategy to be assessed independently. Thus, the strong performance obtained using the same EOM criterion employed in standard HDBSCAN* suggests that much of the gain comes from the graph-topological reformulation itself and demonstrates that the WSS transformation and graph-adapted density formulation produce hierarchies whose stable regions naturally correspond to biologically meaningful flat partitions. This also creates opportunities for future graph-native flat partition-extraction strategies that could further exploit the structural information available in the hierarchy.

An additional strength is the explicit, density-aware identification of uncertain points. In scRNA-seq, such points may repre-sent technical artifacts but also rare or transitional states, so preserving the uncertain set is useful for downstream interpretation and quality control. When a complete partition is required, optional MST-based label propagation (HDBSCAN*(cd,−)) can reassign uncertain points. Although this strategy was originally proposed for semi-supervised settings with externally available labels, here we adapt it to a fully unsupervised setting by using the cluster assignments of other points as labels and propagating them specifically to uncertain points. GraphHDBSCAN* therefore provides three complementary outputs: a family of complete density-based hierarchies spanning multiple *minPts* values, automatically extracted flat partitions retaining explicit uncertain point labels, and optional complete partitions obtained after label propagation. The original uncertain point assignments remain available after propagation, enabling users to distinguish observations assigned directly by the density-based clustering procedure from those assigned during post-processing. Investigating the label-propagated clusters can help characterize specific cell-type populations more precisely and provide insights about the level of heterogeneity within them.

Future work can build on this flexibility through graph-native flat partition extraction, automated graph-construction selection, uncertainty estimates for propagated labels, and integration with multimodal graph representations. The sparse graph formulation and Core-SG acceleration also provide a strong foundation for extending the method to increasingly large single-cell atlases.

Overall, GraphHDBSCAN* unifies similarity graph construction, hierarchical density-based clustering, efficient exploration across multiple *minPts* values, automatic flat partition extraction, explicit uncertain identification, and optional density-aware label propagation within a single framework. The combined hierarchical, flat-partitioning, biological, label-propagation, and runtime results show that GraphHDBSCAN* provides a robust and interpretable approach for recovering both broad and fine-grained structure, including potentially arbitrarily shaped clusters, in high-dimensional single-cell data.

## Methods

### Dataset description

To evaluate the hierarchical clustering, we use three datasets comprising immune and blood cell types, whose annotations were also based on orthogonal information such as physical cell sorting or protein abundance (via CITE-seq protocol). Moreover, as immune cell differentiation processes have been extensively studied, we leverage well-established marker-based cell hierarchies as a ground-truth biological reference to evaluate the performance of the evaluated algorithms.

The dataset introduced by Zheng et al.^36^ consists of reads from fluorescence-activated cell sorting (FACS)-sorted peripheral blood mononuclear cells (PBMCs). The second PBMC dataset, which we refer to as the CITE-seq dataset, is derived from a protocol that enables simultaneous profiling of single-cell transcriptomes and surface protein expression. In this dataset, sorted immune cell populations were labeled using hashtag antibodies, and protein abundances were used to derive ground-truth annotations. Finally, the Bone-marrow dataset provides one of the deepest views of the bone-marrow stem and progenitor compartments to date with up to 85 cell types^39^. In this dataset, we compare hierarchical clustering results to labels that were generated with a combination of protein abundances (CITE-seq), cell sorting and label transfer from a distinct blood-cell atlas (see^39^ for details).

We use seventeen scRNA-seq datasets preprocessed by Szmigiel et al.^12^ to evaluate the performance of our method in flat partitioning settings. These datasets used for quantitative clustering evaluation provide reference cell-type annotations, which were obtained by methods orthogonal to clustering such as cell sorting and cell genotyping, which enables an objective and unbiased evaluation. The preprocessing of the datasets included filtering low-quality cells and lowly expressed genes. Normalization was performed by scaling each cell to have a total count equal to the median total count across all cells, followed by a log(1 + *x*) transformation. Finally, the 1,000 most highly variable genes were selected for downstream analysis and further evaluation.

These include several datasets containing cells from embryonic development, which provide an unbiased ground-truth reference. The dataset by Biase et al.^49^ contains cell-type labels, as they were assigned based on distinct time points during embryonic development. A similar approach was used in the Deng et al.^50^ dataset, where individual cells were dissociated from embryos at various stages of mouse pre-implantation development and labeled accordingly. Likewise, the Goolam dataset^51^ captures cells collected at successive stages of early embryonic development, including the late 2-cell, late 4-cell, late 8-cell, 16-cell, and 32-cell stages. In the Jung dataset^52^, cell populations were isolated using a FACS protocol. The dataset by Koh et al.^53^ also consists of FACS-purified H7 human embryonic stem cells at various differentiation stages. The dataset by Kolodziejczyk et al.^54^ contains mouse embryonic stem cells cultured under three different conditions, forming three distinct cell populations. Another noteworthy dataset is from Yan et al.^55^, which includes 124 human pre-implantation embryos and embryonic stem cells. Because of their origin in well-defined developmental stages, the cell-type annotations in this dataset are considered highly reliable. We also use the benchmark datasets introduced by Tian et al.^47^, where the ground truth is defined based on known genetic differences that identify specific cell lines. Finally, we include liver cells from the first version of the Tabula Sapiens dataset^56^, in which cell-level manual annotation was performed by tissue experts on a two-dimensional UMAP embedding.

For the evaluation of label propagation, we use the PBMC3k dataset, a widely used PBMC scRNA-seq dataset as part of SCANPY tutorials. It was selected because, under this setting, a substantial fraction of cells is initially identified as uncertain, providing a clear example in which many observations require reassignment. Here, we follow a standard `SCANPY`-based preprocessing workflow for PBMC3k^9,36^. After standard quality-control filtering, library-size normalization, log transformation, highly variable gene selection, regression of total counts, and scaling, PCA with 100 components is computed and used to construct the weighted similarity graph supplied to GraphHDBSCAN*. Importantly, the use of PCA here is specific to this dedicated label-propagation example and is neither required by GraphHDBSCAN* nor used to assess its clustering performance. An overview of all datasets and their specific roles in the experimental evaluation is provided in Supplementary Table S1.

### Data representation

Given high-dimensional observations (e.g., scRNA-seq expression profiles), we first construct a *similarity graph* in which nodes represent samples and weighted edges encode local affinities (see Supplementary Material section Similarity Graph). In our experiments, we begin by building a *k*NN graph using either Euclidean or cosine distances and reweighting edges using standard similarity functions, including Gaussian and UMAP kernels, or Jaccard similarity (see Supplementary Material section “Constructing a similarity graph from the data to use as an input for GraphHDBSCAN*”). The resulting weighted similarity graph constitutes the data representation supplied to GraphHDBSCAN*.

### GraphHDBSCAN*: Graph-based Hierarchical Clustering

In this paper, we propose GraphHDBSCAN*, a graph-based hierarchical clustering method that takes a weighted similarity graph as input, computes pairwise weighted structural similarities on its edges, and converts these similarities into dissimilarities. The method then reformulates the principal concepts of HDBSCAN* to operate directly on the resulting sparse, edge-weighted graph.

To efficiently explore multiple *minPts* values, we use Core-SG^24^ machinery. When a flat partition is desired, it is obtained with FOSC/FOSC-X^29,30^, and uncertain points can optionally be assigned to the most suitable clusters by label propagation^31^.

Below, we briefly outline the main components of GraphHDBSCAN*; detailed descriptions are provided in the Supplementary Material section “GraphHDBSCAN*”.

### Weighted Structural Similarity transformation

To enhance robustness and effectiveness in high-dimensional settings, we further compute a WSS, which generalizes SNN similarity^25^ and the structural similarity used in SCAN^57^ to the weighted-graph setting (see Supplementary Material section “Why WSS? Weighted Structural Similarity and Its Relationship to SCAN”). The resulting weighted similarity graph captures both agreement in neighborhoods and the strength of connections. Finally, we transform similarity weights into dissimilarities (*d* = 1 − similarity) so that the graph is compatible with density-based clustering concepts such as core dissimilarity and MRD (Supplementary Material section “Weighted Structural Similarity transformation”).

### Hierarchical density clustering

We then apply the graph-adapted formulation of HDBSCAN* to this graph-based dissimilarity representation. Edge weights are treated as dissimilarities when computing core dissimilarities, MRDs, and the corresponding MST. We adapt the definitions of core distance and MRD to operate on sparse, edge-weighted graphs, ensuring that the resulting algorithm behaves correctly even though the graph does not contain all pairwise dissimilarities (Supplementary Material section “Applying graph-adapted HDBSCAN*”).

To efficiently explore multiple settings of the density-smoothing factor, *minPts*, we employ the Core-SG machinery, which allows all required MSTs to be derived from a single compact spanning graph instead of recomputing MSTs for each *minPts* separately. This enables GraphHDBSCAN* to efficiently compute a broad family of exact hierarchies without requiring the user to provide as input a single *minPts* value before clustering (Supplementary Material section “Core-SG and Its Role in GraphHDBSCAN*”).

### Flat partitioning and label propagation

From the hierarchical clustering obtained by GraphHDBSCAN*, we extract a flat partition using FOSC/FOSC-X with the EOM criterion originally proposed for HDBSCAN*, favoring clusters that persist over a wide range of density levels (Supplementary Material section “Hierarchical and flat clustering outputs”). As in standard density-based clustering, some points are labeled as uncertain. In our scRNA-seq experiments, we optionally reassign these uncertain points to the most appropriate clusters using the MST-based label propagation strategy HDBSCAN*(cd,−)^31^. Although this strategy was originally proposed for semi-supervised settings in which external labels are available for a subset of the data, here we adapt it to a completely unsupervised setting by using the cluster assignments of non-uncertain points as the labeled subset and propagating these labels specifically to observations identified as uncertain. To our knowledge, this is the first use of this strategy for the specific purpose of post-hoc uncertain-point reassignment in a fully unsupervised clustering setting (see Supplementary Material section “HDBSCAN*(cd,−) and Its Use for Uncertain-Point Reassignment”). In this procedure, all non-uncertain samples are treated as labeled, and labels are propagated along MST paths in mutual reachability space, assigning each uncertain point to the densest accessible region. Since the MST is already available from the clustering step, this post-processing does not increase the asymptotic complexity (Supplementary Material section “Label propagation”).

### Identification of cell-surface markers using sc2marker

To identify the cell-surface markers related to each subpopulation identified by the density hierarchy, we use sc2marker^33^ v1.0.3. This tool uses a one-versus-all approach to identify antibodies associated with cluster-specific markers, which can then be used to isolate the cell clusters of interest.

### Understanding GraphHDBSCAN* uncertain points and label propagation

To further understand the characteristics of the uncertain points relabeled by the label propagation approach, we used a gene marker selection approach. To this end, the label-propagated cells were identified using the label-propagated label. Next, a gene marker approach using the `sc.pp.rank_genes` function was performed to select genes (features) specific to each cluster. Our main hypothesis here is that the selected features of the LP clusters are closer to the LP labels than all the rest of the cell clusters. Finally, we compute Pearson correlations between clusters, using the top 20 genes for each cluster (original cell types and LP cell types).

### Benchmarking GraphHDBSCAN* Flat Partitioning

We evaluated all algorithms under two scenarios: *Default*, using their default configurations, and *Best*, involving extensive tuning to select the configuration maximizing ARI for each algorithm and dataset. AMI was reported for the same ARI-selected configuration.

For SC3 and FlowSOM, the known number of reference classes was supplied as the target number of clusters in the Default scenario. In the Best scenario, the number of clusters was optimized, with the true number included among the candidate values. Tuning focused on the number of clusters for SC3, and additionally on distance metrics, self-organizing map dimensions, and training iterations for FlowSOM.

For HDBSCAN* and HDBSCAN*+PCA, tuning focused on *minPts*. The latter was included to assess whether dimensionality reduction could improve HDBSCAN* performance on high-dimensional scRNA-seq data. For both methods, ARI and AMI were calculated after excluding points labeled as noise, and the resulting scores were multiplied by the proportion of non-noise points in the dataset, following Moulavi et al.^58^. This penalizes solutions that leave a large fraction of observations unclustered. For all graph-based methods (GraphHDBSCAN*, Louvain, Leiden, AHSCAN, DHSCAN, and Paris), graph-construction settings were optimized as part of the data-representation pipeline, separately from the clustering algorithms’ intrinsic hyperparameters, although both were considered within the same evaluation procedure. For Louvain and Leiden, all algorithm-specific hyperparameters were comprehensively tuned following Szmigiel et al.^12^. AHSCAN and DHSCAN automatically selected flat partitions by maximizing their respective modularity criteria, whereas Paris required a horizontal cut yielding the known number of reference classes in both scenarios. For GraphHDBSCAN*, the Best scenario selected the highest-performing solution from the family of hierarchies generated over *minPts* values from 2 to 50, efficiently derived from a single initial computation without independently rerunning the clustering procedure for each value. ARI and AMI were computed after optional label propagation, which assigns uncertain points to clusters and yields complete flat partitions.

Complete tuning was feasible across all 17 datasets for HDBSCAN*, HDBSCAN*+PCA, AHSCAN, Paris, and FlowSOM. For DHSCAN, it was computationally infeasible on Zheng, TabulaSapiensLiver_cxg, and CITE-seq, and for SC3 on Tian sce_sc_10x_5cl_qc, Zheng, TabulaSapiensLiver_cxg, and CITE-seq. In these cases, the Default configurations were retained.

## Supporting information

Supplementary_Data_sc2marker_Complete_results

## Acknowledgements

This project has been funded by the Novo Nordisk Foundation (#NNF23OC0079660), the Federal Ministry of Research, Technology and Space of Germany (BMFTR) via the Consortia E:MED Fibromap and CureFib for I.G.C. and the German Research Foundation (DFG; 3888802535, CRU344-417911533) for I.G.C..

## Author contributions statement

S.A.G. and R.J.G.B.C. conceived the work and designed the algorithm. S.A.G. implemented the code based on the proposed method and produced all the software libraries and documentation for its public distribution. S.A.G., A.W.S. and J.N. performed the experiments and contributed to writing the manuscript. S.A.G., A.W.S., J.N., I.G.C. and R.J.G.B.C. analyzed and interpreted the results. A.Z., R.J.G.B.C. and I.G.C. reviewed and edited the manuscript. R.J.G.B.C. and I.G.C. supervised the project. R.J.G.B.C. secured the funding.

## Additional information

### Accession codes

The package and examples demonstrating how to use it to reproduce the results are available at the GraphHDBSCAN* repository.

## Supplementary Material

### Background

#### Related Work

Graph clustering encompasses a broad range of methods developed for different objectives and data representations. Here, we restrict the discussion to methods most directly related to GraphHDBSCAN*: (i) density-based graph-clustering methods that extend concepts such as core points and density connectivity to network data, and (ii) graph-clustering methods that explicitly produce hierarchical cluster structures. We additionally consider methods operating on sparse similarity graphs constructed from feature-vector data when their setting is particularly close to ours. Broader classes of flat community-detection and representation-learning methods fall outside the scope of this discussion.

SCAN^57^ is a graph-based counterpart of DBSCAN. On unweighted graphs, it defines structural similarity from neighborhood overlap and identifies core vertices according to two global hyperparameters: a structural-similarity threshold *ε* and a minimum structural-neighborhood size *µ*. As in DBSCAN, the use of a single global threshold can be restrictive when clusters exhibit different levels of structural density, and the method produces a flat clustering for a selected pair of hyperparameter values. SCAN additionally identifies vertices that do not belong to clusters as hubs or outliers. SCOT+HintClus^59^ builds on this structural clustering formulation using ideas related to OPTICS^60^ to expose clustering structure across different similarity levels. However, its representation is based on an ordering/reachability view rather than the explicit density-based hierarchy and automatic multi-level cluster extraction provided by HDBSCAN*-style methods.

AHSCAN^13^ and DHSCAN^14^ explicitly construct hierarchical clusterings of graph data. AHSCAN follows an agglomerative strategy, whereas DHSCAN follows a divisive strategy based on structural similarity. Their original formulations operate on unweighted graphs, so differences in edge strengths are not incorporated into the structural-similarity measure. Moreover, when a flat partition is required, a single level of the hierarchy is selected using a modularity-based criterion. Such a horizontal selection applies a common level of granularity across the hierarchy and does not provide the explicit density-based noise treatment or local multi-level cluster selection available in HDBSCAN*-style hierarchies.

Paris^17^ instead operates directly on weighted graphs and constructs a hyperparameter-free agglomerative hierarchy using a node-pair-sampling-based distance. It therefore preserves edge-weight information and provides a multiscale dendrogram. However, its hierarchy is based on successive agglomerative merges rather than density connectivity, all observations remain assigned within the hierarchy, and obtaining a flat partition requires an additional dendrogram-selection step.

More recently, hierarchy-oriented methods have also been proposed specifically for high-dimensional and single-cell data, including scMustree^18^ and Bonsai^19^. scMustree constructs terminal cell populations through iterative PCA, *k*-nearest-neighbor (kNN) graph construction, and community detection, and subsequently organizes these populations into a hierarchy through bottom-up merging guided by cluster-specific functional rankings. Bonsai instead reconstructs a maximum-likelihood tree under a probabilistic model of high-dimensional observations and their measurement uncertainty. These methods therefore pursue different methodological objectives: scMustree constructs a functional hierarchy from recursively derived terminal cell populations, whereas Bonsai performs probabilistic tree reconstruction in a high-dimensional continuous space. In contrast, GraphHDBSCAN* constructs a density-based hierarchy directly from a sparse weighted similarity graph, explicitly models density connectivity, and retains low-density observations as uncertain. scMustree and Bonsai are nevertheless included in our empirical evaluation because both provide hierarchical representations of high-dimensional single-cell data and therefore constitute relevant contemporary alternatives for comparison.

GraphHDBSCAN* therefore lies at the intersection of graph-based and density-based hierarchical clustering. In contrast to methods designed primarily for native network data, our setting starts from a sparse weighted similarity graph representing high-dimensional feature-vector observations. GraphHDBSCAN* then defines density and density connectivity directly on this graph and produces a family of density-based hierarchies, while preserving the possibility of extracting non-horizontal flat clusterings and explicit uncertain (noise) assignments.

#### Sparse Similarity Graphs

Graph-based clustering of feature-vector data commonly begins by constructing a sparse similarity graph *G* = (*V, E, w*), in which each vertex *v_i_* ∈ *V* represents an observation *x_i_* and weighted edges encode selected local relationships between observations. Rather than retaining all pairwise relationships, a graph construction policy determines which proximities are represented by edges; pairs that are not retained are equivalently assigned zero similarity and are absent from the sparse graph. Different policies can be used to determine the edge set, including *k*NN, mutual-*k*NN, and threshold-based neighborhood graphs. Once the graph topology has been determined, its edges can additionally be weighted or reweighted using a similarity function. Consequently, graph construction involves conceptually distinct choices concerning which local relationships are retained and how the strengths of those retained relationships are represented.

These choices determine properties such as graph sparsity, connectivity, and local neighborhood structure and can therefore affect the behavior of downstream graph-clustering algorithms^12^. Importantly, however, the construction of this representation is separate from the clustering algorithm itself. GraphHDBSCAN*, like other graph-based clustering methods, receives the resulting sparse weighted similarity graph as input. The particular graph constructions used in our experiments are described in the Data Representation section below.

#### HDBSCAN*

HDBSCAN*^15,16^ is a hierarchical extension of DBSCAN^27^ designed to identify clusters across different density levels while allowing observations in low-density regions to be identified as noise. Unlike DBSCAN, which requires the specification of a single global density threshold, HDBSCAN* constructs a complete density-based hierarchy by varying this threshold.

The only compulsory hyperparameter is *minPts*, which is inherited from DBSCAN. For each observation *x_i_*, the *core distance* is defined as

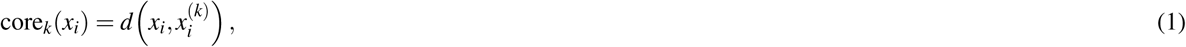

where *x*^(*k*)^_*i*_ is the *k*-th nearest neighbor of *x_i_*, with *k* := *minPts*. The core distance has a direct density interpretation: it is the smallest neighborhood radius for which *x_i_*contains the minimum local mass required by *minPts*. Equivalently, its inverse is associated with the largest density level at which the observation can still satisfy this local-density requirement.

Using the core distances, HDBSCAN* defines the *mutual reachability distance* between two observations *x_i_* and *x _j_* as

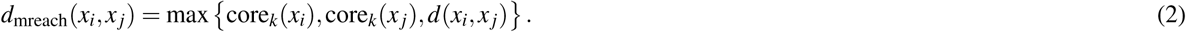

This transformation combines local density and direct pairwise connectivity. At any distance (i.e., neighborhood radius) threshold *ε* corresponding to a density threshold 1*/ε*, two observations can be both dense and directly density connected if and only if their pairwise distance and both of their core distances do not exceed *ε*. Thus, edge weights in the transformed space of a conceptual, fully-connected mutual-reachability graph reflect the local density and density-connectivity requirements imposed by the corresponding weights as variable density thresholds according to a non-parametric density estimate with density smoothing factor given by *minPts*.

An MST of such a conceptual mutual-reachability graph provides a compact representation from which the complete hierarchy can be recovered. Starting from the lowest density level and progressively increasing the density threshold (equivalently, decreasing *ε*), edges are removed from the MST and connected components evolve accordingly. Importantly, this process does not consist only of successive cluster splits. Many edge removals cause peripheral observations to leave clusters and become noise, while only particular edge removals produce substantive splits into multiple clusters. The result is therefore a density-based hierarchy containing both nested clusters and explicit noise assignments.

The hierarchy may optionally be further simplified by imposing a minimum cluster size, *min_cluster_size*. This is an optional additional condensation of the hierarchy rather than a requirement for constructing the density hierarchy itself. Flat clusters can subsequently be selected from the hierarchy using stability-based criteria such as the Excess-of-Mass criterion employed within FOSC/FOSC-X^29,30^.

#### Core-SG and Its Role in GraphHDBSCAN*

While HDBSCAN* effectively builds hierarchical density-based cluster structures through the computation of an MST in the space of MRDs, exploring multiple values of the density-smoothing hyperparameter *minPts* can be computationally expensive. In a straightforward implementation, a separate density-based MST is computed for each value of *minPts*, requiring the corresponding MRDs to be recomputed for each setting. The CORE-distance based Spanning Graph (Core-SG) framework^24^ addresses the cost of these repeated computations by showing that, after an initial MST is obtained for the largest *minPts* value under consideration, the exact MSTs associated with smaller *minPts* values can be recovered from a substantially sparser replacement graph.

More specifically, for a range of *minPts* ∈ [1*, k*_max_], Core-SG shows that all the corresponding MSTs can be exactly derived from the union of the following two graph structures:

1. the MST computed using the largest considered *minPts* value *k*_max_, denoted MST*_k_*_max_ , and
2. the *k*_max_-nearest-neighbor graph (*k*_max_-NNG), including all ties at the *k*_max_-th nearest neighbor. Accordingly, the CORE-distance based Spanning Graph is defined as

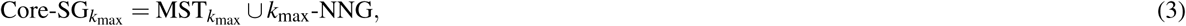

and acts as a replacement graph containing the edges required to derive the MSTs associated with smaller *minPts* values. The theoretical justification is given by the inclusion relation

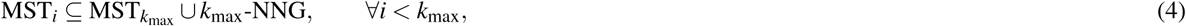

which guarantees that the MST corresponding to each smaller density-smoothing setting can be recovered from Core-SG*_k_*_max_ without repeating the original complete-graph computation.

Importantly, Core-SG does not reduce the cost of computing the initial MST*_k_*_max_ . In the standard complete-graph formulation, this initial computation still involves a graph with *O*(*n*^2^) pairwise relationships. The computational advantage arises *after* this initial step: assuming *k*_max_ ≪ *n*, Core-SG*_k_*_max_ contains only *O*(*nk*_max_) edges, and the MSTs associated with the remaining *minPts* values can therefore be computed from this sparse replacement graph rather than from the original complete graph.

An additional optimization, denoted Core-SG*, further reduces redundancy by iteratively removing edges that do not belong to any MST within the range of *minPts* values under consideration. This produces a minimal replacement graph preserving the edges required to recover the corresponding MSTs exactly. Consequently, after the initial computation at *k*_max_, Core-SG and Core-SG* enable the exact MSTs associated with multiple density-smoothing settings to be obtained with substantially lower additional computational and memory requirements than independently recomputing each MST.

In GraphHDBSCAN*, we exploit this same theoretical machinery in the graph-based setting. An initial hierarchy is computed for the largest considered value *k*_max_, after which the exact MSTs, and consequently the corresponding clustering hierarchies, for smaller *minPts* values can be efficiently derived. GraphHDBSCAN* can therefore return a family of exact density-based hierarchies spanning multiple density-smoothing settings without requiring the user to select and tune a single *minPts* value before clustering.

### HDBSCAN*(cd,−) and Its Use for Uncertain-Point Reassignment

HDBSCAN*(cd,−) was originally introduced as a semi-supervised classification strategy within the unified density-based framework by Gertrudes et al.^31^. In its original setting, class labels are externally available for a subset of the observations, and the density structure encoded by the mutual-reachability minimum spanning tree (MST) is used to propagate those labels to unlabeled observations.

In this work, we use the same strategy for a different purpose. Rather than relying on externally provided class labels, we use the cluster labels obtained by GraphHDBSCAN* itself: observations assigned to clusters in the extracted flat partition are treated as labeled, whereas observations identified as uncertain are treated as unlabeled. The HDBSCAN*(cd,−) procedure is then used specifically to reassign these uncertain observations to the most appropriate clusters. Therefore, the overall clustering procedure remains completely unsupervised. To the best of our knowledge, this is the first use of HDBSCAN*(cd,−) for post-hoc reassignment of uncertain observations using cluster labels generated internally by the clustering algorithm.

### Label propagation via density connectivity

Let *X_L_* denote the observations assigned to clusters by GraphHDBSCAN*, together with their corresponding cluster labels *Y_L_*, and let *X_U_* denote the observations identified as uncertain. HDBSCAN*(cd,−) uses the MST of the mutual-reachability graph as the backbone for propagation. Each uncertain observation is assigned the label of the labeled observation that can be reached through the path minimizing the maximum mutual-reachability distance:

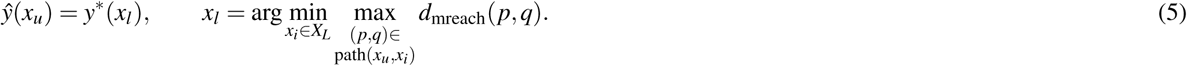

The minimax path criterion favors assignments through the strongest density-connected path, which can be proven to be always contained within the paths along the density-based MST, rather than simply assigning an uncertain observation to the nearest cluster according to the original pairwise distance. Because the MST required by this procedure has already been obtained during GraphHDBSCAN*, the label-propagation step reuses the existing hierarchical clustering structure.

### Data representation

#### Constructing a similarity graph from the data to use as an input for GraphHDBSCAN*

As illustrated in Fig. 1a, the data representation stage is separate from GraphHDBSCAN* and produces the sparse weighted similarity graph supplied to the clustering algorithm. This representation is not specific to GraphHDBSCAN*: the same type of graph representation is also required when graph-based clustering methods such as Louvain or Leiden are applied to feature-vector data.

In our experiments, we first construct an initial *k*NN graph directly from the feature-vector data using either Euclidean or cosine distance. To distinguish this (data representation related) neighborhood-size hyperparameter from the variable density-smoothing factor *minPts* used later by GraphHDBSCAN*, we denote the former by *k*_NN_. Notice that the distance measure used at this stage determines which observations are retained as local neighbors and should therefore be distinguished from the subsequent edge-weighting strategy.

Once the neighborhood topology has been constructed, the retained edges are assigned or reweighted using one of the graph-weighting strategies considered in our experiments: Gaussian, UMAP-based, or Jaccard/PhenoGraph weighting. These strategies determine the strength of the local relationships represented by the edges of the graph, whereas Euclidean or cosine distance is used for the preceding nearest-neighbor search. The resulting sparse weighted similarity graph constitutes the input to GraphHDBSCAN*. In particular, when Jaccard weighting is used, the resulting Jaccard values are simply the edge weights of the input graph; the WSS transformation is subsequently applied internally by GraphHDBSCAN* to this weighted graph.

A further practical consideration is graph connectivity. In order to construct a single MST spanning all observations, and therefore a complete clustering hierarchy, the graph processed by GraphHDBSCAN* must be connected. When the sparse similarity graph is disconnected, we consider two alternatives. The first is to return to the data-representation stage and increase *k*_NN_ until a connected similarity graph is obtained. This retains the interpretation of all graph edges as relationships derived directly from the original data, at the cost of a denser graph. Alternatively, the disconnected components can be retained and subsequently joined by edges assigned the maximum possible dissimilarity. This preserves the internal hierarchy of each component while connecting the components only at the highest dissimilarity level, producing a coarser top-level hierarchy.

### GraphHDBSCAN*

GraphHDBSCAN* receives a sparse weighted similarity graph as input and defines a density-based hierarchical clustering formulation directly on this graph. As shown in Fig. 1a, the method first uses WSS to represent the similarity between adjacent vertices in terms of their weighted neighborhood structure. It then defines graph-based counterparts of the HDBSCAN* core dissimilarity and MRD directly on this sparse representation, from which the corresponding minimum spanning trees and density-based clustering hierarchies are obtained. Thus, GraphHDBSCAN* is not an application of conventional HDBSCAN* to a transformed feature space; its density and connectivity concepts are reformulated to operate on a sparse weighted graph. The proposed GraphHDBSCAN* algorithm (Algorithm 1) consists of the following steps:

1. Weighted Structural Similarity transformation. Compute pairwise weighted structural similarities on the edges of the input similarity graph and assign the resulting WSS values as edge weights. These similarities are then transformed into dissimilarities to ensure consistency with the concepts of core distance and MRD.
2. Applying graph-adapted HDBSCAN*. Use the transformed edge dissimilarities to define graph-based core dissimilarities and MRDs directly from the sparse WSS dissimilarity graph. These quantities are used to construct the corresponding density-based MST and recover the complete density-based clustering hierarchy. Core-SG^24^ is used to efficiently derive the exact MSTs, and consequently the corresponding hierarchies, associated with a range of *minPts* values from the initial computation at the largest considered value.
3. Hierarchical and flat clustering outputs. Produce a family of full density-based clustering hierarchies, one for each considered value of *minPts*. Each hierarchy can be represented as a condensed tree for visualization and cluster extraction. When a flat partition is desired, clusters are selected from a chosen hierarchy using FOSC/FOSC-X with the EOM criterion^29,30^, while observations not assigned to any selected cluster remain uncertain.
4. Optional label propagation. Observations identified as uncertain in the flat partition can optionally be assigned to the most suitable clusters using the semi-supervised HDBSCAN*(cd,−) procedure^31^.

In the following sections, we describe these steps in detail.

### Weighted Structural Similarity transformation

One effective strategy for addressing the challenges associated with the curse of dimensionality is employing techniques like SNN^25^. In this paper, we utilize a WSS^21^ graph as an advanced form of SNN, explicitly incorporating similarity weights between nodes.

WSS constitutes the first stage of GraphHDBSCAN*. It receives the weighted similarity graph constructed during the separate data representation stage and computes pairwise weighted structural similarity to reweight the edges of the input similarity graph. The weighted structural similarity^21^ measure *σ* (*u, v*) between two nodes *u* and *v* in a graph is defined as:

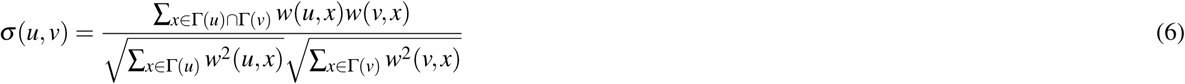

where:

- *σ* (*u, v*) is the similarity score between nodes *u* and *v*.
- Γ(*u*) and Γ(*v*) are the sets of neighbors of nodes *u* and *v*, respectively.
- *w*(*u, x*) and *w*(*v, x*) are the weights of the edges between nodes *u* and *x*, and *v* and *x*, respectively.

The WSS measure can be interpreted as the cosine similarity between the weighted neighborhood vectors associated with *u* and *v*. More precisely, each vertex is represented by a vector whose entries are the weights of its connections to neighboring vertices, with zero entries for vertices outside its neighborhood. The numerator in Eq. (6) is therefore the inner product of these two vectors, for which only shared neighbors contribute non-zero terms, while the denominator normalizes by their Euclidean norms.

Consequently, WSS measures the agreement between the complete weighted neighborhood profiles of two vertices. Two adjacent vertices receive a high WSS value when their connections are concentrated on the same neighbors with similar relative strengths, whereas differences in either neighborhood membership or edge-weight profiles decrease their structural similarity. In the GraphHDBSCAN* implementation, we additionally provide optional choices concerning how the WSS graph is constructed. In particular, users may choose whether each vertex is included in its own neighborhood for the WSS computation and whether WSS is evaluated only for pairs already connected in the input similarity graph or also for previously non-adjacent pairs having non-zero WSS.

For all experiments in this paper, however, we use a single fixed configuration. Following the usual sparse SNN setting, WSS is computed only for edges already present in the input similarity graph, so the WSS transformation reweights the existing graph without introducing new connections. We also include each vertex itself in its neighborhood. This distinguishes an existing edge whose two endpoints otherwise have no common neighbors from the absence of an edge between two vertices.

Finally, because the graph-adapted density formulation below is expressed in terms of dissimilarities, WSS is converted to a structural dissimilarity by

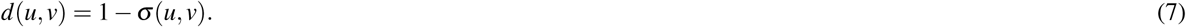

Since *σ* (*u, v*) ∈ [0, 1], the resulting dissimilarities also lie in [0, 1], with larger values corresponding to weaker structural agreement.

### Why WSS? Weighted Structural Similarity and Its Relationship to SCAN

The motivation for using WSS in GraphHDBSCAN* is to obtain a local structural similarity that exploits the graph representation of the data rather than relying directly on the original pairwise edge weights. Starting from the sparse weighted similarity graph supplied as input to GraphHDBSCAN*, WSS reweights its edges by combining agreement between local neighborhoods with the strengths of the corresponding connections. In this way, WSS incorporates both shared-neighborhood information, which can be advantageous in high-dimensional settings, and the graded similarity information already encoded by the input graph. The relationship between WSS and SCAN provides an additional interpretation of this transformation. SCAN^57^ uses structural similarity based on neighborhood overlap in unweighted graphs. WSS generalizes this structural similarity to weighted graphs by additionally accounting for the strengths of the connections to shared neighbors. Importantly, this relationship does not mean that GraphHDBSCAN* is simply an extension of the SCAN algorithm. Rather, WSS provides the structural similarity representation on which the graph-adapted density concepts of GraphHDBSCAN*, including core dissimilarity and MRD, are subsequently defined.

As shown below, when the input graph is binary and self-loops are included, WSS reduces exactly to the structural similarity used by SCAN. This special case also allows us to establish a more precise relationship between the core-cluster structures recovered by SCAN and GraphHDBSCAN*.

### Reduction to SCAN’s structural similarity in the binary case

Now, consider the special case where the graph is unweighted, i.e., all edge weights are either 0 or 1, and where self-loops are added so that every node includes itself in its neighborhood. In this 0/1 setting:

- The numerator becomes the size of the shared neighborhood between *u* and *v*:

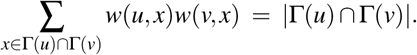

- The denominator becomes the geometric mean of the neighborhood sizes:

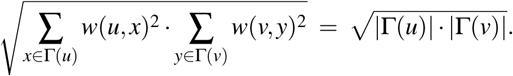

Therefore, in the 0/1 case with self-loops, the WSS expression simplifies to:

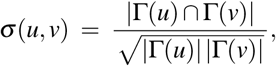

which is precisely the structural similarity measure originally used in SCAN. That is, WSS reduces exactly to SCAN’s similarity in the binary case.

### Equivalence of SCAN and GraphHDBSCAN* When Using WSS on Binary Graphs

We now show the detailed connection between SCAN and GraphHDBSCAN* when WSS is used on a binary graph. For corresponding values of SCAN’s *µ* and GraphHDBSCAN*’s *minPts* under the same neighborhood-counting convention, GraphHDBSCAN* with MRD based on WSS recovers the same core-cluster structure that SCAN produces for varying thresholds *ε*, disregarding the assignment of the so-called *border* points.

First we should recall the following about SCAN:

1. SCAN declares *u* and *v* to be neighbors if *σ* (*u, v*) ≥ *ε*.
2. A node is a “core” if it satisfies the minimum structural-neighborhood requirement defined by *µ*.
3. SCAN then merges nodes via *ε*-reachability (transitive closure through core nodes and high-similarity edges).

On the GraphHDBSCAN* side, let *k* = *minPts* and define

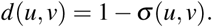

Following the standard HDBSCAN* definition, let

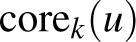

denote the distance from *u* to its *k*th nearest neighbor according to *d*, under the same neighborhood-counting convention used to match SCAN’s *µ*. The corresponding mutual-reachability distance is

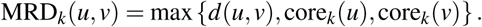

For corresponding choices of *µ* and *k* = *minPts* that impose the same core-neighborhood requirement, a node *u* is a core node in SCAN at threshold *ε* if and only if

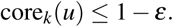

Now let

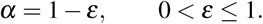

For two core nodes *u* and *v*,

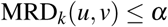

if and only if:

- *σ* (*u, v*) ≥ 1 − *α* = *ε*,
- *u* satisfies the SCAN core condition at threshold *ε*,
- and *v* does as well.

Therefore, at the corresponding levels *α* = 1 − *ε*, the graph induced by core vertices and edges satisfying MRD(*u, v*) ≤ *α* has the same connected components as the graph formed by structurally similar core vertices in SCAN. Since thresholding the MST preserves the connected components of the corresponding thresholded mutual-reachability graph, GraphHDBSCAN* and

SCAN recover the same core-cluster structure at the corresponding thresholds.

This equivalence concerns the core-cluster structure. SCAN may additionally assign non-core vertices to clusters through direct structural reachability from core vertices. These border-point assignments are therefore not included in the equivalence.

We see that our proposed GraphHDBSCAN*, which is equipped with WSS, is a strict generalization of SCAN with its structural similarity:

- On general weighted graphs, it respects both neighborhood structure and similarity strength.
- On binary graphs, WSS reduces exactly to SCAN’s structural similarity.
- For corresponding values of SCAN’s *µ* and GraphHDBSCAN*’s *minPts*, GraphHDBSCAN* recovers the same corecluster structure as SCAN across similarity thresholds, with *ε* = 1 − *α*.

In short, GraphHDBSCAN* with WSS on a binary graph provides a full hierarchical representation of SCAN’s core-cluster structure across similarity thresholds. Whereas SCAN requires a particular similarity threshold *ε* to be selected in order to obtain a corresponding flat clustering, GraphHDBSCAN* automatically represents the core-cluster structures associated with the full range of corresponding thresholds within a single hierarchy.

#### Applying graph-adapted HDBSCAN*

The second stage of GraphHDBSCAN* reformulates the principal concepts of HDBSCAN* so that they operate directly on the sparse, edge-weighted dissimilarity graph produced by the WSS transformation. In standard HDBSCAN*, core distances and mutual reachability distances are defined from pairwise dissimilarities in the original feature space. Here, the corresponding concepts are defined from the structural dissimilarities represented by the edges of the WSS graph.

Let

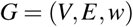

denote the WSS dissimilarity graph, where *V* is the vertex set, *E* ⊆ *V* × *V* is the set of retained graph edges, and *w* : *E* → [0, 1] denotes their structural dissimilarities. The bounded range follows directly from the WSS construction: *σ* (*u, v*) ∈ [0, 1] and *w*(*u, v*) = 1 − *σ* (*u, v*).

To avoid confusion with the neighborhood-size hyperparameter *k*_NN_ used during the separate similarity-graph construction stage, we denote the GraphHDBSCAN* density-smoothing factor by *m* := *minPts* in the definitions below.

**Core dissimilarity.** For each vertex *v* ∈ *V* , let

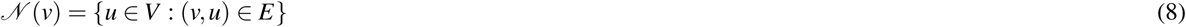

be the set of vertices connected to *v*, and let

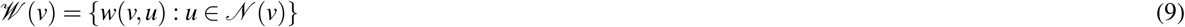

be the corresponding incident structural dissimilarities. Let *w*(m)(*v*) denote the *m*-th order statistic of *W* (*v*), i.e., its *m*-th smallest value. We define the core dissimilarity of *v* as

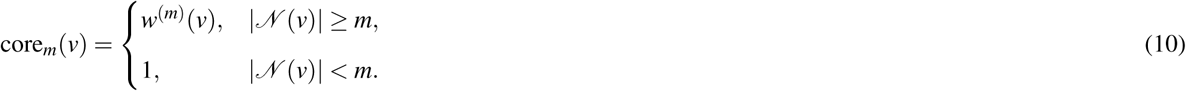

This definition preserves the density interpretation of the ordinary HDBSCAN* core distance in the graph setting. For any structural dissimilarity threshold *ε <* 1, define

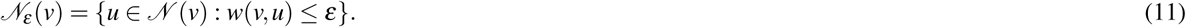

Then, whenever *v* has at least *m* represented neighbors, core*_m_*(*v*) is precisely the smallest value of *ε* for which

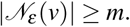

Thus, analogously to HDBSCAN*, the core dissimilarity is the smallest structural-neighborhood radius at which the vertex contains the local mass required by *minPts*. Equivalently, under the usual *λ* = 1*/ε* interpretation, its inverse determines the largest density level at which the vertex can still satisfy this local-density requirement. If fewer than *m* neighbors are represented in the sparse graph, no threshold *ε <* 1 can satisfy this condition, and the core dissimilarity is assigned the maximum value 1.

### Extended structural dissimilarity

Because the input graph is sparse, most vertex pairs have no explicitly represented edge. We therefore define

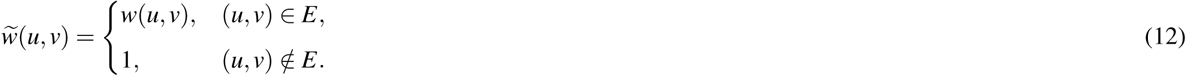

The value 1 assigned to a missing edge should not be interpreted as an observed pairwise dissimilarity. Rather, it represents the absence of a direct structural relationship in the sparse input graph.

### Mutual reachability dissimilarity

The graph-based MRD is then defined as

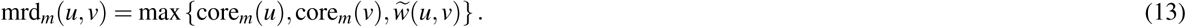

By construction, mrd*_m_*(*u, v*) ∈ [0, 1].

This definition also retains the density-connectivity interpretation of the ordinary HDBSCAN* mutual reachability distance. For any *ε <* 1,

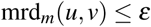

only if (i) *u* and *v* are directly connected in the WSS graph with structural dissimilarity at most *ε*, and (ii) both vertices have at least *m* represented neighbors within their respective structural *ε*-neighborhoods. Mutual reachability therefore combines two requirements: sufficient local structural density around both endpoints and structural connectivity between them. As *ε* decreases, or equivalently as the density level *λ* = 1*/ε* increases, weak structural connections and vertices that no longer satisfy the local-density requirement are progressively removed, yielding the density-based hierarchy.

**Interpretation of** mrd*m* = 1. Since each node is included in its own neighborhood, any existing edge (*u, v*) ∈ *E* guarantees that Γ(*u*) ∩Γ(*v*) ̸= 0/ and therefore the WSS similarity *σ* (*u, v*) in Eq. 6 is strictly positive. As a result, the corresponding dissimilarity *d*(*u, v*) = 1 − *σ* (*u, v*) always satisfies *d*(*u, v*) ∈ [0, 1) for every edge in the graph. Hence, within the original WSS dissimilarity graph, no edge ever attains a dissimilarity value of 1. When we assign a dissimilarity value of 1 to non-adjacent vertex pairs (*u, v*) ∈*/E* in Eq. 12, this value therefore serves as an explicit indicator of absence of structural evidence rather than an empirical large distance.

Furthermore, from the core dissimilarity definition in Eq. 10, a vertex with fewer than *m* incident edges also receives a core dissimilarity of 1. Consequently, from Eq. 13, we obtain

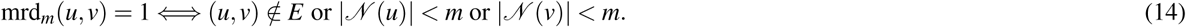

Thus, the value mrd*_m_*= 1 consistently denotes a lack of structural support: either the two vertices are not directly connected in the WSS graph, or at least one endpoint cannot serve as an *m*-core point due to insufficient local connectivity.

**Compatibility with Core-SG.** Core-SG was originally formulated for the conventional HDBSCAN* setting, where the underlying pairwise dissimilarity graph can be viewed as complete. In GraphHDBSCAN*, however, the WSS representation is already sparse. The definitions above allow the Core-SG machinery to be used without constructing an additional complete graph.

Conceptually, Eq. (12) completes the sparse graph by assigning dissimilarity 1 to all missing pairs. In practice, this completion does not need to be materialized. Provided that the WSS graph is connected and that every vertex has at least *k*_max_ represented neighbors, where *k*_max_ is the largest *minPts* value in the Core-SG range, all required core dissimilarities are strictly smaller than 1. Moreover, because the connected WSS graph contains a spanning tree consisting of represented edges with structural dissimilarities below 1, missing edges with dissimilarity 1 are not required to construct the relevant MSTs.

Under the same condition, the *k*_max_-nearest-neighbor graph required by Core-SG is obtained from the represented structural dissimilarities and is itself a subgraph of the already sparse WSS graph. Consequently, the Core-SG replacement graph is effectively a sparse graph extracted from the sparse WSS representation, rather than a second graph requiring construction of the implicit complete graph. No modification of the Core-SG inclusion principle is required; the graph-based definitions of core dissimilarity and MRD above provide the quantities on which the same MST-recovery machinery operates.

For the *k*NN-based input representations used in our experiments, this condition can be ensured by choosing the graph neighborhood size such that every vertex retains at least *k*_max_ neighbors. If this condition is not satisfied, the GraphHDBSCAN* definitions remain well-defined through Eqs. (10)–(13), but the sparse Core-SG construction no longer provides the same computational advantage; in that case, the input representation can instead be reconstructed with a larger neighborhood size.

#### Hierarchical and flat clustering outputs

The primary output of GraphHDBSCAN* is a family of full density-based clustering hierarchies, one for each considered value of the density-smoothing factor *minPts*. Each hierarchy is obtained from its corresponding mutual-reachability MST. As the dissimilarity threshold is progressively decreased (equivalently, as the density level increases), MST edges are removed and the connected components evolve.

Importantly, this process should not be interpreted simply as a conventional dendrogram in which every edge removal necessarily splits one cluster into two new subclusters. Many edge removals simply cause peripheral observations to detach from a cluster and become uncertain points. Only a comparatively small subset of edge removals corresponds to genuine split events in which one cluster separates into two or more density-connected subclusters. Consequently, the hierarchy naturally represents both the nested organization of clusters and the progressive identification of uncertain points as the density level increases.

This unique characteristic can be harnessed for the purposes of visualization and subsequent cluster extraction, by simplifying the full hierarchy and representing it instead as a so-called condensed tree, which emphasizes the relevant cluster-splitting structure while retaining the information about observations that leave clusters as uncertain observations. Further controlled condensation can be optionally imposed through a (non-compulsory) *min_cluster_size* hyperparameter. Values greater than one suppress branches corresponding to clusters smaller than the specified size and therefore further condense the hierarchy. Importantly, *min_cluster_size* is not required for constructing the density-based hierarchy; it is an optional post-processing choice that controls the degree of additional condensation, being particularly useful for exploratory visualization purposes.

When a non-hierarchical representation is desired, a flat partition can be extracted from a selected hierarchy. GraphHDB-SCAN* does not require a horizontal cut at a single global density level. Instead, we use the FOSC/FOSC-X framework^29,30^ with the EOM criterion to select a non-overlapping collection of clusters from different parts and potentially different density levels of the hierarchy. This allows clusters supported at different depths along different branches to coexist in the same flat solution.

Observations that do not belong to any of the selected clusters retain their explicit uncertain labels. Thus, for any selected hierarchy, GraphHDBSCAN* can provide both the original density-based flat partition containing uncertain observations and, if desired, a complete partition in which those uncertain observations are subsequently reassigned through the optional label-propagation procedure described below.

#### Label propagation

Label propagation constitutes an optional post-processing stage applied to a flat clustering solution and does not modify the hierarchical outputs of GraphHDBSCAN*. Density-based clustering naturally leaves some observations labeled as uncertain. When a complete partition is desired, we use HDBSCAN*(cd,−)^31^ to reassign these observations according to the density-connectivity structure already recovered by GraphHDBSCAN*.

HDBSCAN*(cd,−) was originally proposed for semi-supervised classification, where class labels are externally available for a subset of the observations and are propagated to the remaining unlabeled observations through a mutual-reachability MST. Our use of this procedure is different and remains completely unsupervised. For a flat partition extracted from a GraphHDBSCAN* hierarchy, observations assigned directly to clusters are treated as labeled using their internally generated cluster labels, while observations identified as uncertain are treated as unlabeled. The HDBSCAN*(cd,−) propagation rule is then used specifically to assign those uncertain observations to the most density-connected labeled clusters. To the best of our knowledge, this constitutes the first use of this semi-supervised strategy for the specific purpose of reassigning uncertain observations in a completely unsupervised clustering setting.

Importantly, this post-processing step does not require GraphHDBSCAN* to select an additional value of *minPts* or to construct a new MST. As described above, Core-SG provides the collection of exact MSTs associated with the considered range of *minPts* values. For a flat partition extracted from a particular hierarchy, label propagation reuses the corresponding already-computed MST. Consequently, the additional post-processing cost is associated with the label-propagation procedure itself, rather than with recomputing mutual-reachability MSTs and density-based hierarchies.

For an unlabeled uncertain observation *x_u_*, HDBSCAN*(cd,−) considers paths in the corresponding mutual-reachability MST leading to labeled observations. The preferred assignment is determined by the labeled observation reachable through the path minimizing its maximum mutual-reachability dissimilarity. Thus, labels are propagated according to density connectivity rather than simply according to the original pairwise proximity, favoring assignments that avoid crossing weak density connections between clusters.

The procedure is optional: the original uncertain labels produced by GraphHDBSCAN* are always retained alongside the propagated labels. Users can therefore distinguish observations assigned directly by the density-based clustering procedure from observations assigned only during post-processing and can use either the uncertain-preserving or the complete flat partition depending on the application.

### Runtime analysis

Fig. S3 compares the runtime of Louvain, Leiden, and GraphHDBSCAN* across 17 datasets containing between 49 and 8,508 cells. The datasets are ordered by increasing cell count, and execution time is shown on a logarithmic scale. Runtime includes graph construction from `adata.X`. For each method and dataset, one warm-up run was excluded and five subsequent runs were measured.

Runtime generally increases with dataset size for all three methods. On the smaller and medium-sized datasets, Louvain typically has the lowest median runtime, GraphHDBSCAN* generally lies between Louvain and Leiden, and Leiden is usually the slowest. However, this ordering changes for several of the larger datasets. GraphHDBSCAN* has the lowest median runtime on Tian 10x 5CL, Tabula Sapiens liver, and T-cell CITE-seq. On the largest dataset, Zheng, the runtimes of the three methods are similar.

An important distinction is that a single GraphHDBSCAN* execution provides a family of density-based hierarchies over the considered range of *minPts* values. The input similarity graph and its WSS transformation are computed once. Core-SG then exploits the MST associated with the largest considered value, together with the corresponding sparse neighborhood structure, to recover the exact MSTs for the remaining *minPts* values. Thus, GraphHDBSCAN* does not independently recompute the MST of the implicit mutual-reachability graph from scratch for every density-smoothing setting.

This distinction is relevant when comparing GraphHDBSCAN* with methods such as Louvain and Leiden, which return a single flat community partition for a given setting. GraphHDBSCAN* incurs the additional work required to construct and retain hierarchical clustering information, but a single fitted model supports exploration of multiple density-smoothing settings without repeating the complete data-representation and clustering procedure independently for each *minPts* value. Overall, no method is consistently the fastest across all datasets. GraphHDBSCAN* remains competitive throughout the benchmark and becomes the fastest method on several of the larger datasets, indicating practical runtime scaling over the evaluated range.

**Algorithm 1.**
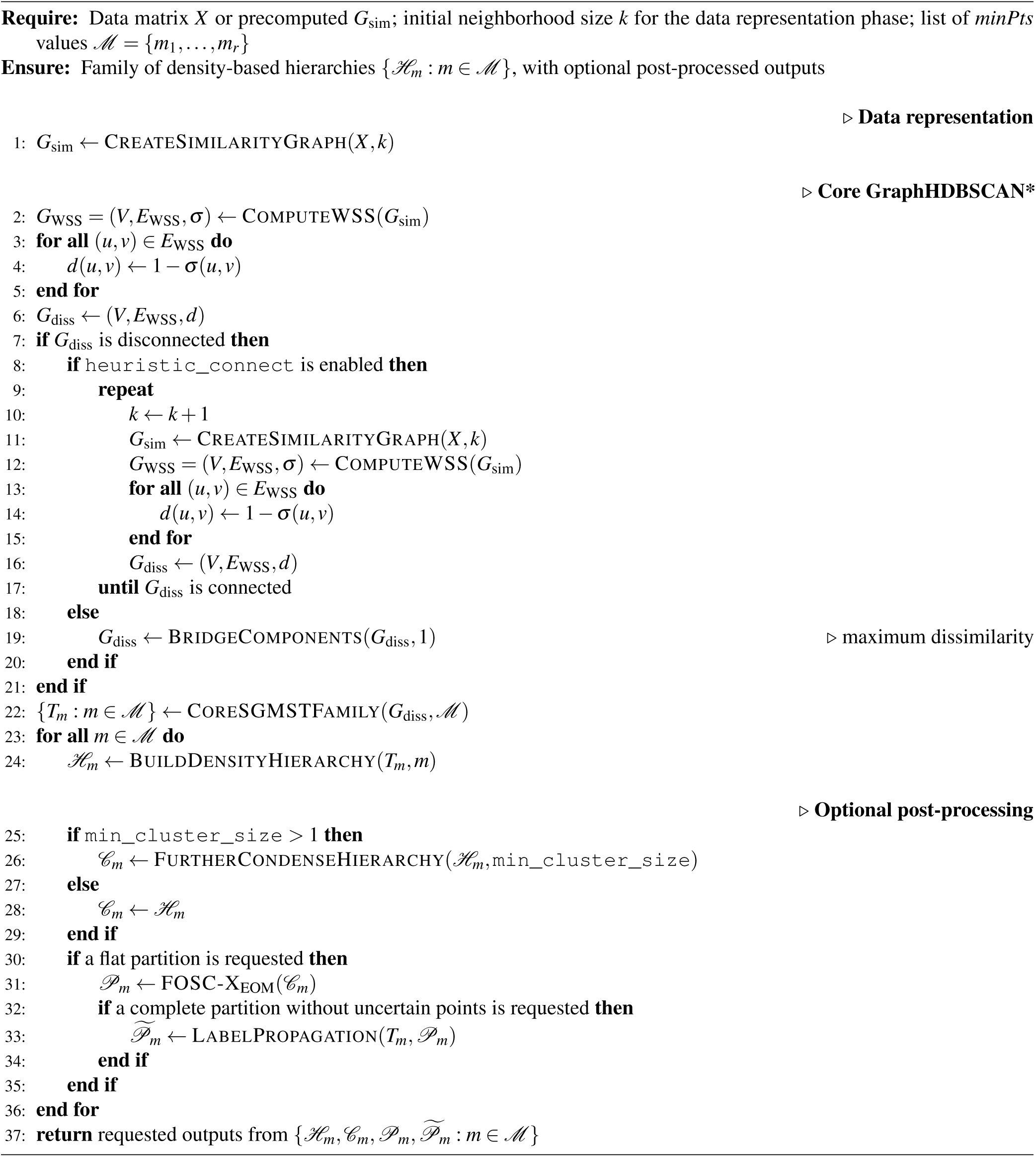
GraphHDBSCAN* workflow.

|  |  |  |
|---|---|---|
| **Require:** | Data matrix *X* or precomputed *G*<sub>sim</sub>; initial neighborhood size *k* for the data representation phase; list of *minPts* values ℳ = {*m*<sub>1</sub>, …, *m*<sub>*r*</sub>} |  |
| **Ensure:** | Family of density-based hierarchies {ℋ<sub>*m*</sub>: *m* ∈ ℳ}, with optional post-processed outputs |  |
|  |  | ▷ **Data representation** |
| 1: | *G*<sub>sim</sub> ← CreateSimilarityGraph(*X*, *k*) |  |
|  |  | ▷ **Core GraphHDBSCAN\*** |
| 2: | *G*<sub>WSS</sub> = (*V*, *E*<sub>WSS</sub>, σ) ← ComputeWSS(*G*<sub>sim</sub>) |  |
| 3: | **for all** (*u*, *v*) ∈ *E*<sub>WSS</sub> **do** |  |
| 4: | *d*(*u*, *v*) ← 1 − σ(*u*, *v*) |  |
| 5: | **end for** |  |
| 6: | *G*<sub>diss</sub> ← (*V*, *E*<sub>WSS</sub>, *d*) |  |
| 7: | **if** *G*<sub>diss</sub> is disconnected **then** |  |
| 8: | **if** heuristic_connect is enabled **then** |  |
| 9: | **repeat** |  |
| 10: | *k* ← *k* + 1 |  |
| 11: | *G*<sub>sim</sub> ← CreateSimilarityGraph(*X*, *k*) |  |
| 12: | *G*<sub>WSS</sub> = (*V*, *E*<sub>WSS</sub>, σ) ← ComputeWSS(*G*<sub>sim</sub>) |  |
| 13: | **for all** (*u*, *v*) ∈ *E*<sub>WSS</sub> **do** |  |
| 14: | *d*(*u*, *v*) ← 1 − σ(*u*, *v*) |  |
| 15: | **end for** |  |
| 16: | *G*<sub>diss</sub> ← (*V*, *E*<sub>WSS</sub>, *d*) |  |
| 17: | **until** *G*<sub>diss</sub> is connected |  |
| 18: | **else** |  |
| 19: | *G*<sub>diss</sub> ← BridgeComponents(*G*<sub>diss</sub>, 1) | ▷ maximum dissimilarity |
| 20: | **end if** |  |
| 21: | **end if** |  |
| 22: | {*T*<sub>*m*</sub>: *m* ∈ ℳ} ← CoreSGMSTFamily(*G*<sub>diss</sub>, ℳ) |  |
| 23: | **for all** *m* ∈ ℳ **do** |  |
| 24: | ℋ<sub>*m*</sub> ← BuildDensityHierarchy(*T*<sub>*m*</sub>, *m*) |  |
|  |  | ▷ **Optional post-processing** |
| 25: | **if** min_cluster_size > 1 **then** |  |
| 26: | 𝒞<sub>*m*</sub> ← FurtherCondenseHierarchy(ℋ<sub>*m*</sub>, min_cluster_size) |  |
| 27: | **else** |  |
| 28: | 𝒞<sub>*m*</sub> ← ℋ<sub>*m*</sub> |  |
| 29: | **end if** |  |
| 30: | **if** a flat partition is requested **then** |  |
| 31: | 𝒫<sub>*m*</sub> ← FOSC-X<sub>EOM</sub>(𝒞<sub>*m*</sub>) |  |
| 32: | **if** a complete partition without uncertain points is requested **then** |  |
| 33: | 𝒫̃<sub>*m*</sub> ← LabelPropagation(*T*<sub>*m*</sub>, 𝒫<sub>*m*</sub>) |  |
| 34: | **end if** |  |
| 35: | **end if** |  |
| 36: | **end for** |  |
| 37: | **return** requested outputs from {ℋ<sub>*m*</sub>, 𝒞<sub>*m*</sub>, 𝒫<sub>*m*</sub>, 𝒫̃<sub>*m*</sub>: *m* ∈ ℳ} |  |

## Supplementary Tables

**Table S1.** Overview of the datasets used in the experimental evaluation.

| Dataset | Reference labels | Flat partition | Label propagation | Dedicated LP case | Hierarchical clustering | Reference hierarchy | Experimental role and rationale |
|---|---|---|---|---|---|---|---|
| Biase | ✓ | ✓ | ✓ |  |  |  | Contains cells collected at different time points of early embryonic development. |
| Deng | ✓ | ✓ | ✓ |  |  |  | Contains individual cells dissociated from embryos at different stages of mouse pre-implantation development. |
| Goolam | ✓ | ✓ | ✓ |  |  |  | Contains cells collected at the late 2-cell, late 4-cell, late 8-cell, 16-cell, and 32-cell stages. |
| Jung | ✓ | ✓ | ✓ |  |  |  | Contains four cell populations purified using fluorescence-activated cell sorting. |
| Koh | ✓ | ✓ | ✓ |  |  |  | Contains FACS-purified H7 human embryonic stem cells sampled at different differentiation stages. |
| Tian sc_Celseq2_5cl_p1 | ✓ | ✓ | ✓ |  |  |  | A controlled mixture containing 297 cells from five cell lines, measured using UMI counts. The reference labels correspond to cell-line identities determined from known genetic differences rather than from a clustering algorithm. |
| Tian sc_Celseq2_5cl_p2 | ✓ | ✓ | ✓ |  |  |  | A controlled mixture containing 307 cells from five cell lines, measured using UMI counts. Its genetically determined cell-line identities provide reliable reference labels for quantitative evaluation. Together with p1 and p3, it allows the method to be assessed across closely related mixture-control experiments. |
| Tian sc_Celseq2_5cl_p3 | ✓ | ✓ | ✓ |  |  |  | A controlled mixture containing 305 cells from five cell lines, measured using UMI counts. |
| Tian sce_sc_10x_5cl_qc | ✓ | ✓ | ✓ |  |  |  | A controlled mixture containing 3,750 cells from five cell lines, generated using the 10x protocol and represented by UMI counts. It is the largest Tian dataset in this collection and therefore evaluates the method on a higher-throughput dataset while retaining reliable, genetically determined reference labels. |
| Tian sce_sc_CELseq2_qc | ✓ | ✓ | ✓ |  |  |  | A controlled mixture containing 274 cells from three cell lines, generated using the CEL-seq2 protocol and represented by UMI counts. Its reference labels are based on genetically determined cell-line identities. |
| Tian sce_sc_Dropseq_qc | ✓ | ✓ | ✓ |  |  |  | A controlled mixture containing 225 cells from three cell lines, generated using the Drop-seq protocol and represented by UMI counts. It provides genetically determined reference labels and is included to evaluate clustering and label propagation under a different experimental protocol. |
| Tian sce_sc_10x_qc | ✓ | ✓ | ✓ |  |  |  | A controlled mixture containing 902 cells from three cell lines, generated using the 10x protocol and represented by UMI counts. |
| Yan | ✓ | ✓ | ✓ |  |  |  | Contains human pre-implantation embryos and embryonic stem cells. The reference populations correspond to different developmental stages, providing highly confident labels for evaluating whether the recovered flat partition agrees with known human developmental populations. |
| Kolodziejczyk | ✓ | ✓ | ✓ |  |  |  | Contains 704 mouse embryonic stem cells cultured under three different conditions, defining three distinct cell populations. |
| TabulaSapiensLiver_cxg | ✓ | ✓ | ✓ |  |  |  | Contains 5,007 human liver cells assigned to 13 cell types through manual annotation by tissue experts using the CELLxGENE browser and a two-dimensional UMAP representation. |
| Zheng | ✓ | ✓ | ✓ |  | ✓ | ✓ | Contains a large collection of FACS-sorted peripheral blood mononuclear cells representing nine immune-cell populations. Its independently determined labels and large sample size make it suitable for quantitative flat partitioning evaluation. It is also used for hierarchical evaluation because the annotated immune populations have known biological relationships that can be compared with the recovered cluster hierarchy. |
| CITE-seq | ✓ | ✓ | ✓ |  | ✓ | ✓ | Used for both hierarchical clustering and flat partitioning evaluation because its immune-cell annotations were generated using surface-protein measurements, orthogonal to annotations based on RNA-sequencing data. It is used for hierarchical evaluation because the annotated immune populations have known biological relationships that can be compared with the recovered cluster hierarchy. |
| Bone-marrow |  |  |  |  | ✓ | ✓ | One of the largest and most comprehensive datasets representing different lineages of cells in the bone-marrow. Its CITE-seq design pairs gene expression with a panel of surface protein markers, enabling a refined, fine-grained annotation built from multiple clustering solutions and retaining only stable, reproducible cell states. It is used for hierarchical evaluation because of the presence of three-level ground-truth annotation in the dataset. |
| PBMC3k |  |  | ✓ | ✓ |  |  | Used exclusively as the illustrative label propagation case study. It does not provide external ground-truth labels suitable for objective quantitative clustering evaluation, and many cells are initially classified as uncertain. It therefore provides a clear example for demonstrating how label propagation assigns uncertain points to existing clusters while preserving biologically coherent cell populations. |

**Table S2.**
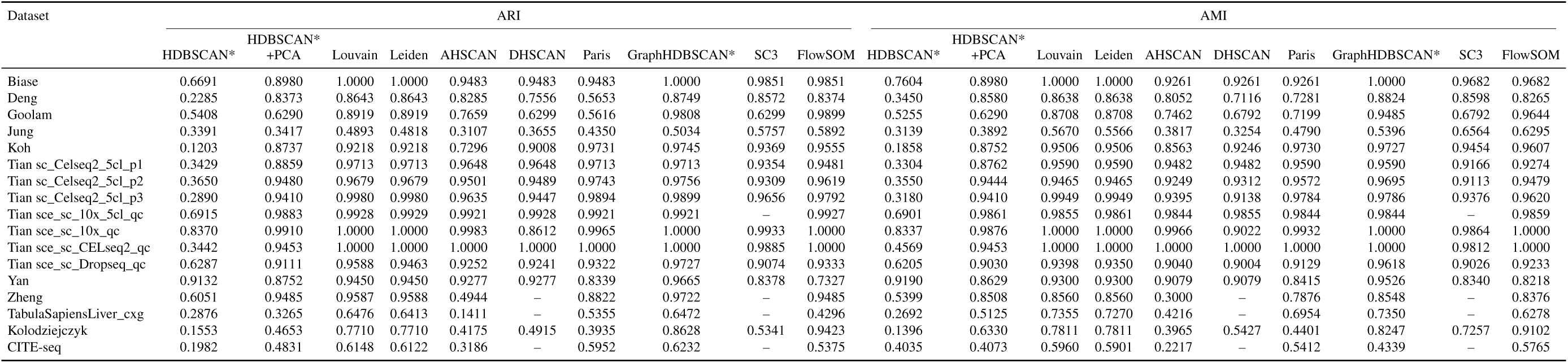
Best-scenario ARI and AMI results for the evaluated clustering methods across the 17 benchmark datasets. Entries marked “–” indicate cases in which a tuned result could not be obtained because tuning was computationally infeasible due to excessive runtime on a particular dataset. For the Best-scenario aggregate analyses, including the boxplots and ranking-based statistical comparisons, the corresponding Default-scenario result was retained in these cases.

| Dataset | ARI |  |  |  |  |  |  |  |  |  | AMI |  |  |  |  |  |  |  |  |  |
|---|---|---|---|---|---|---|---|---|---|---|---|---|---|---|---|---|---|---|---|---|
|  | HDBSCAN* | HDBSCAN*<br>+PCA | Louvain | Leiden | AHSCAN | DHSCAN | Paris | GraphHDBSCAN* | SC3 | FlowSOM | HDBSCAN* | HDBSCAN*<br>+PCA | Louvain | Leiden | AHSCAN | DHSCAN | Paris | GraphHDBSCAN* | SC3 | FlowSOM |
| Biase | 0.6691 | 0.8980 | 1.0000 | 1.0000 | 0.9483 | 0.9483 | 0.9483 | 1.0000 | 0.9851 | 0.9851 | 0.7604 | 0.8980 | 1.0000 | 1.0000 | 0.9261 | 0.9261 | 0.9261 | 1.0000 | 0.9682 | 0.9682 |
| Deng | 0.2285 | 0.8373 | 0.8643 | 0.8643 | 0.8285 | 0.7556 | 0.5653 | 0.8749 | 0.8572 | 0.8374 | 0.3450 | 0.8580 | 0.8638 | 0.8638 | 0.8052 | 0.7116 | 0.7281 | 0.8824 | 0.8598 | 0.8265 |
| Goolam | 0.5408 | 0.6290 | 0.8919 | 0.8919 | 0.7659 | 0.6299 | 0.5616 | 0.9808 | 0.6299 | 0.9899 | 0.5255 | 0.6290 | 0.8708 | 0.8708 | 0.7462 | 0.6792 | 0.7199 | 0.9485 | 0.6792 | 0.9644 |
| Jung | 0.3391 | 0.3417 | 0.4893 | 0.4818 | 0.3107 | 0.3655 | 0.4350 | 0.5034 | 0.5757 | 0.5892 | 0.3139 | 0.3892 | 0.5670 | 0.5566 | 0.3817 | 0.3254 | 0.4790 | 0.5396 | 0.6564 | 0.6295 |
| Koh | 0.1203 | 0.8737 | 0.9218 | 0.9218 | 0.7296 | 0.9008 | 0.9731 | 0.9745 | 0.9369 | 0.9555 | 0.1858 | 0.8752 | 0.9506 | 0.9506 | 0.8563 | 0.9246 | 0.9730 | 0.9727 | 0.9454 | 0.9607 |
| Tian sc_Celseq2_5cl_p1 | 0.3429 | 0.8859 | 0.9713 | 0.9713 | 0.9648 | 0.9648 | 0.9713 | 0.9713 | 0.9354 | 0.9481 | 0.3304 | 0.8762 | 0.9590 | 0.9590 | 0.9482 | 0.9482 | 0.9590 | 0.9590 | 0.9166 | 0.9274 |
| Tian sc_Celseq2_5cl_p2 | 0.3650 | 0.9480 | 0.9679 | 0.9679 | 0.9501 | 0.9489 | 0.9743 | 0.9756 | 0.9309 | 0.9619 | 0.3550 | 0.9444 | 0.9465 | 0.9465 | 0.9249 | 0.9312 | 0.9572 | 0.9695 | 0.9113 | 0.9479 |
| Tian sc_Celseq2_5cl_p3 | 0.2890 | 0.9410 | 0.9980 | 0.9980 | 0.9635 | 0.9447 | 0.9894 | 0.9899 | 0.9656 | 0.9792 | 0.3180 | 0.9410 | 0.9949 | 0.9949 | 0.9395 | 0.9138 | 0.9784 | 0.9786 | 0.9376 | 0.9620 |
| Tian sce_sc_10x_5cl_qc | 0.6915 | 0.9883 | 0.9928 | 0.9929 | 0.9921 | 0.9928 | 0.9921 | 0.9921 | – | 0.9927 | 0.6901 | 0.9861 | 0.9855 | 0.9861 | 0.9844 | 0.9855 | 0.9844 | 0.9844 | – | 0.9859 |
| Tian sce_sc_10x_qc | 0.8370 | 0.9910 | 1.0000 | 1.0000 | 0.9983 | 0.8612 | 0.9965 | 1.0000 | 0.9933 | 1.0000 | 0.8337 | 0.9876 | 1.0000 | 1.0000 | 0.9966 | 0.9022 | 0.9932 | 1.0000 | 0.9864 | 1.0000 |
| Tian sce_sc_CELseq2_qc | 0.3442 | 0.9453 | 1.0000 | 1.0000 | 1.0000 | 1.0000 | 1.0000 | 1.0000 | 0.9885 | 1.0000 | 0.4569 | 0.9453 | 1.0000 | 1.0000 | 1.0000 | 1.0000 | 1.0000 | 1.0000 | 0.9812 | 1.0000 |
| Tian sce_sc_Dropseq_qc | 0.6287 | 0.9111 | 0.9588 | 0.9463 | 0.9252 | 0.9241 | 0.9322 | 0.9727 | 0.9074 | 0.9333 | 0.6205 | 0.9030 | 0.9398 | 0.9350 | 0.9040 | 0.9004 | 0.9129 | 0.9618 | 0.9026 | 0.9233 |
| Yan | 0.9132 | 0.8752 | 0.9450 | 0.9450 | 0.9277 | 0.9277 | 0.8339 | 0.9665 | 0.8378 | 0.7327 | 0.9190 | 0.8629 | 0.9300 | 0.9300 | 0.9079 | 0.9079 | 0.8415 | 0.9526 | 0.8340 | 0.8218 |
| Zheng | 0.6051 | 0.9485 | 0.9587 | 0.9588 | 0.4944 | – | 0.8822 | 0.9722 | – | 0.9485 | 0.5399 | 0.8508 | 0.8560 | 0.8560 | 0.3000 | – | 0.7876 | 0.8548 | – | 0.8376 |
| TabulaSapiensLiver_cxg | 0.2876 | 0.3265 | 0.6476 | 0.6413 | 0.1411 | – | 0.5355 | 0.6472 | – | 0.4296 | 0.2692 | 0.5125 | 0.7355 | 0.7270 | 0.4216 | – | 0.6954 | 0.7350 | – | 0.6278 |
| Kolodziejczyk | 0.1553 | 0.4653 | 0.7710 | 0.7710 | 0.4175 | 0.4915 | 0.3935 | 0.8628 | 0.5341 | 0.9423 | 0.1396 | 0.6330 | 0.7811 | 0.7811 | 0.3965 | 0.5427 | 0.4401 | 0.8247 | 0.7257 | 0.9102 |
| CITE-seq | 0.1982 | 0.4831 | 0.6148 | 0.6122 | 0.3186 | – | 0.5952 | 0.6232 | – | 0.5375 | 0.4035 | 0.4073 | 0.5960 | 0.5901 | 0.2217 | – | 0.5412 | 0.4339 | – | 0.5765 |

**Table S3.** Assigning cell-type identities to clusters (PBMC3k tutorial)

| Cluster ID | Markers | Cell Type |
|---|---|---|
| 0 | IL7R, CCR7 | Naive CD4<sup>+</sup> T cells |
| 1 | CD14, LYZ | CD14<sup>+</sup> monocytes |
| 2 | IL7R, S100A4 | Memory CD4<sup>+</sup> T cells |
| 3 | MS4A1 | B cells |
| 4 | CD8A | CD8<sup>+</sup> T cells |
| 5 | FCGR3A, MS4A7 | FCGR3A<sup>+</sup> monocytes |
| 6 | GNLY, NKG7 | NK cells |
| 7 | FCER1A, CST3 | Dendritic cells |
| 8 | PPBP | Platelets |

## Supplementary Figures

**Figure S1.**
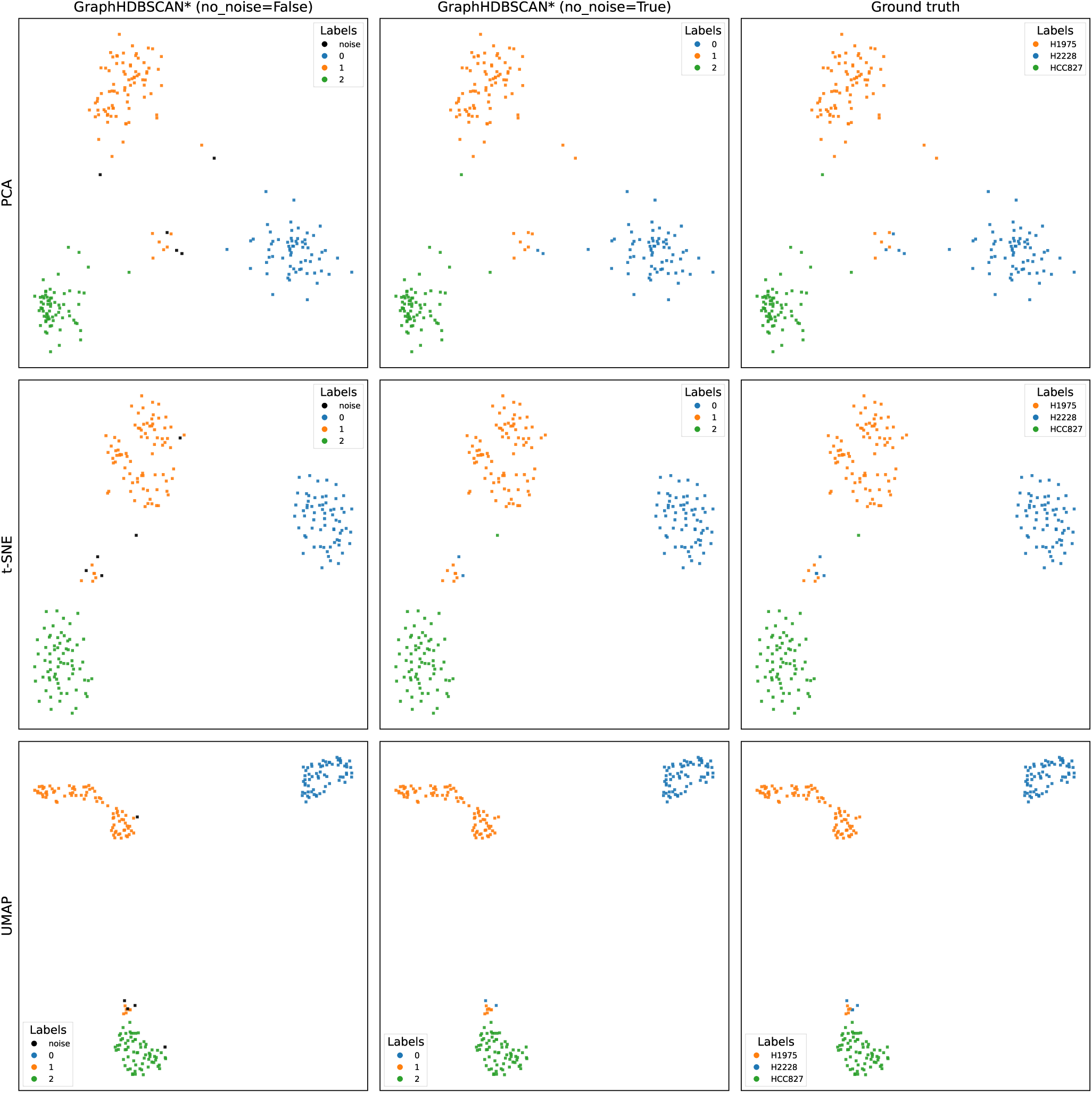
Visualization of GraphHDBSCAN* before and after label propagation on the Tian sce_sc_Dropseq_qc dataset. Rows show PCA, t-SNE, and UMAP representations, respectively. The left column shows the GraphHDBSCAN* partition before label propagation, including points identified as uncertain; the middle column shows the partition after label propagation; and the right column shows the reference cell-line labels (H1975, H2228, and HCC827). The comparison illustrates how uncertain points are reassigned to clusters that visually agree with the corresponding reference populations across the three low-dimensional representations.

**Figure S2.**
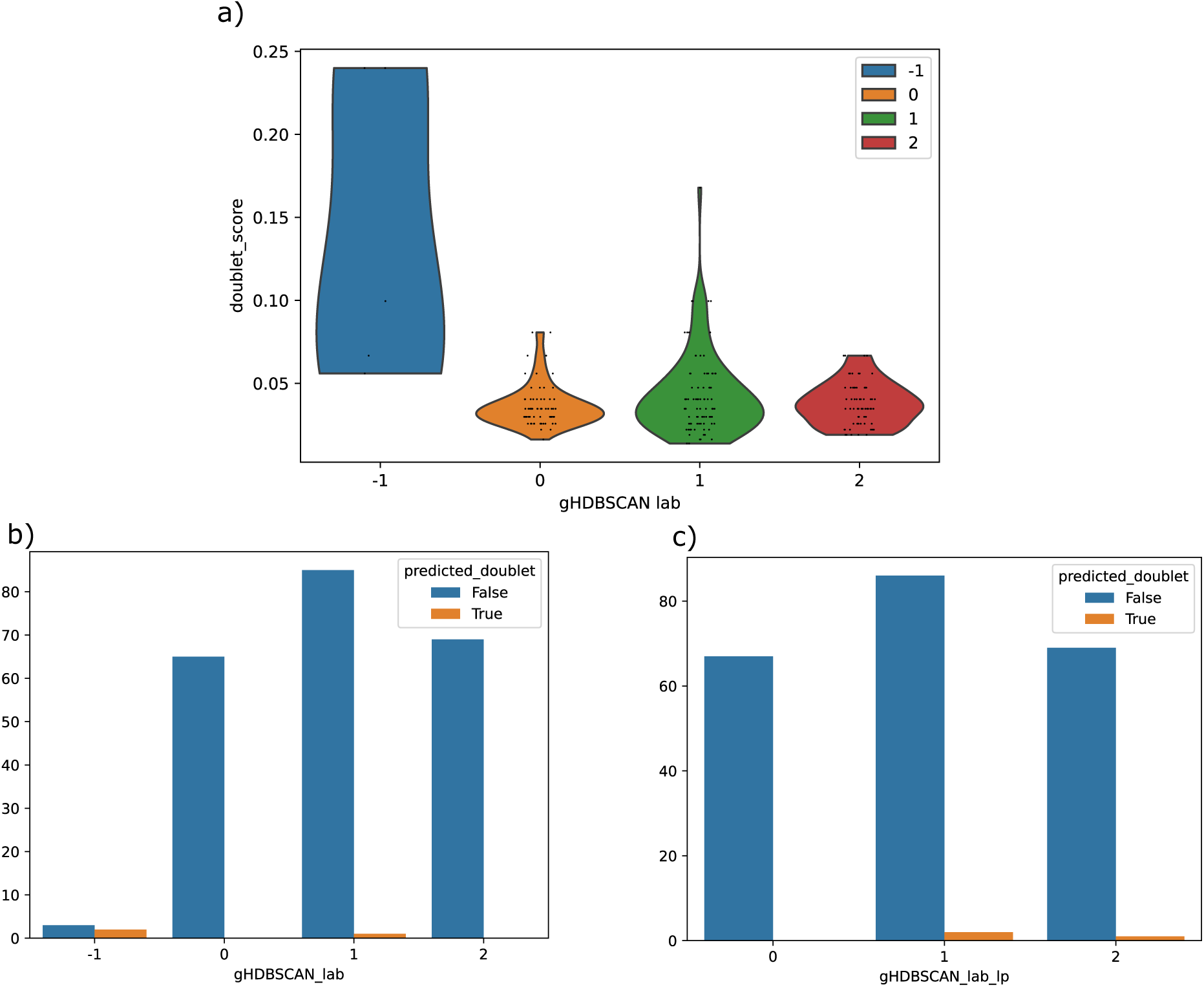
Doublet analysis for the Tian sce_sc_Dropseq_qc dataset. (a) Doublet-score distribution across GraphHDBSCAN* clusters before label propagation, including the points identified as uncertain. (b–c) Doublet predictions per cluster before and after label propagation.

**Figure S3.**
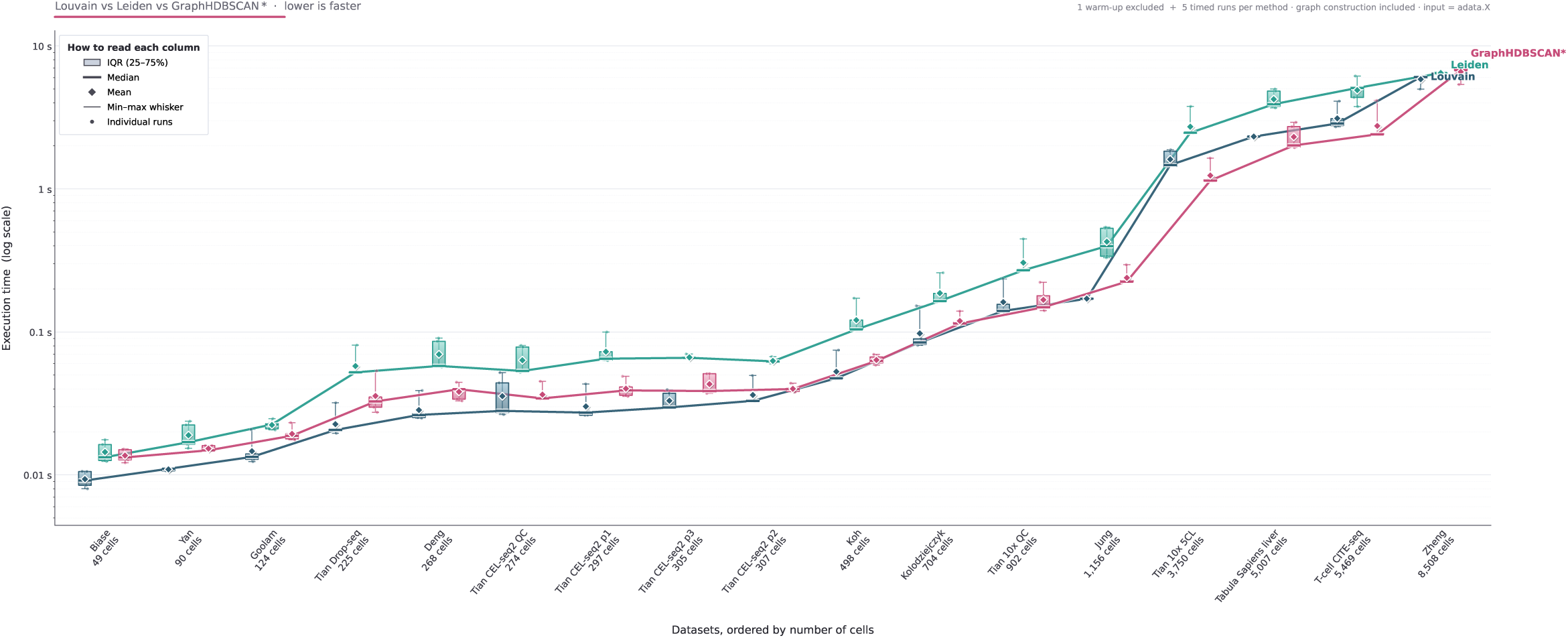
Runtime comparison of Louvain, Leiden, and GraphHDBSCAN* across datasets ordered by increasing cell count. Runtime includes graph construction, with one warm-up run excluded and five timed runs recorded per method and dataset. Boxes show the interquartile range, center lines indicate medians, diamonds indicate means, whiskers show the minimum and maximum, and points represent individual runs. Connected lines trace median runtime across datasets. Execution time is shown on a logarithmic scale.

**Figure S4.**
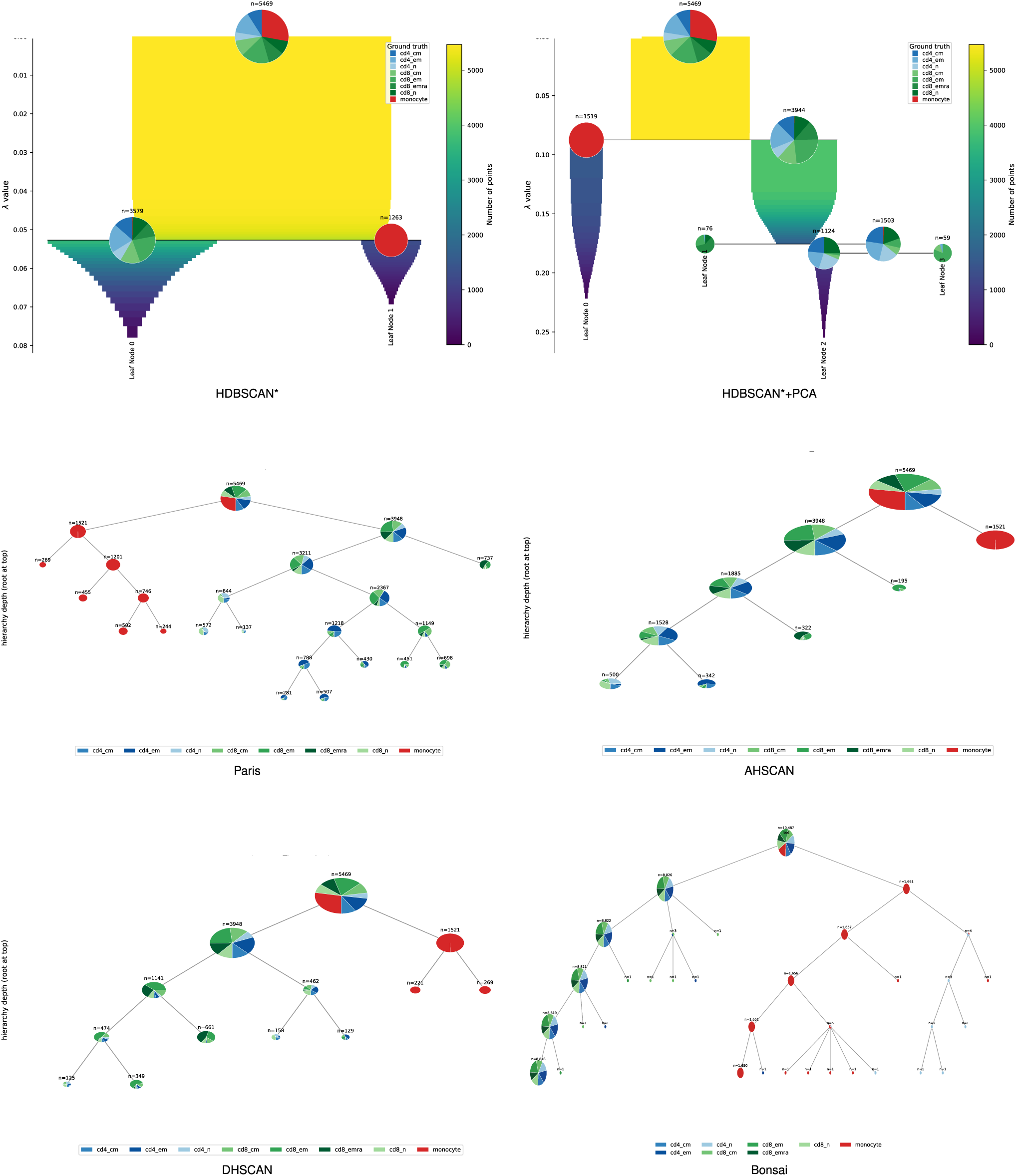
Hierarchical clustering results for the CITE-seq dataset obtained with HDBSCAN*, HDBSCAN*+PCA, Paris, AHSCAN, DHSCAN, and Bonsai. The vertical axes represent method-specific hierarchy statistics that are not comparable across methods. Pie charts show cluster composition according to the biological reference labels (red for monocytes; blue colors for CD4 T cells and green for CD8 T cells). HDBSCAN* produces a very coarse hierarchy, while HDBSCAN*+PCA introduces considerably more structure. Most methods can find the separation of monocytes (red) from T-cell populations (green and blue), but several internal branches remain mixed and the finer CD4/CD8 organization is only partially preserved, with the exception of Paris and HDBSCAN*+PCA, which can split some CD4 from some CD8 populations, as reflected by their higher HAI scores in contrast to other competing methods. Bonsai detects very small clusters and cannot split T-cell subtypes.

**Figure S5.**
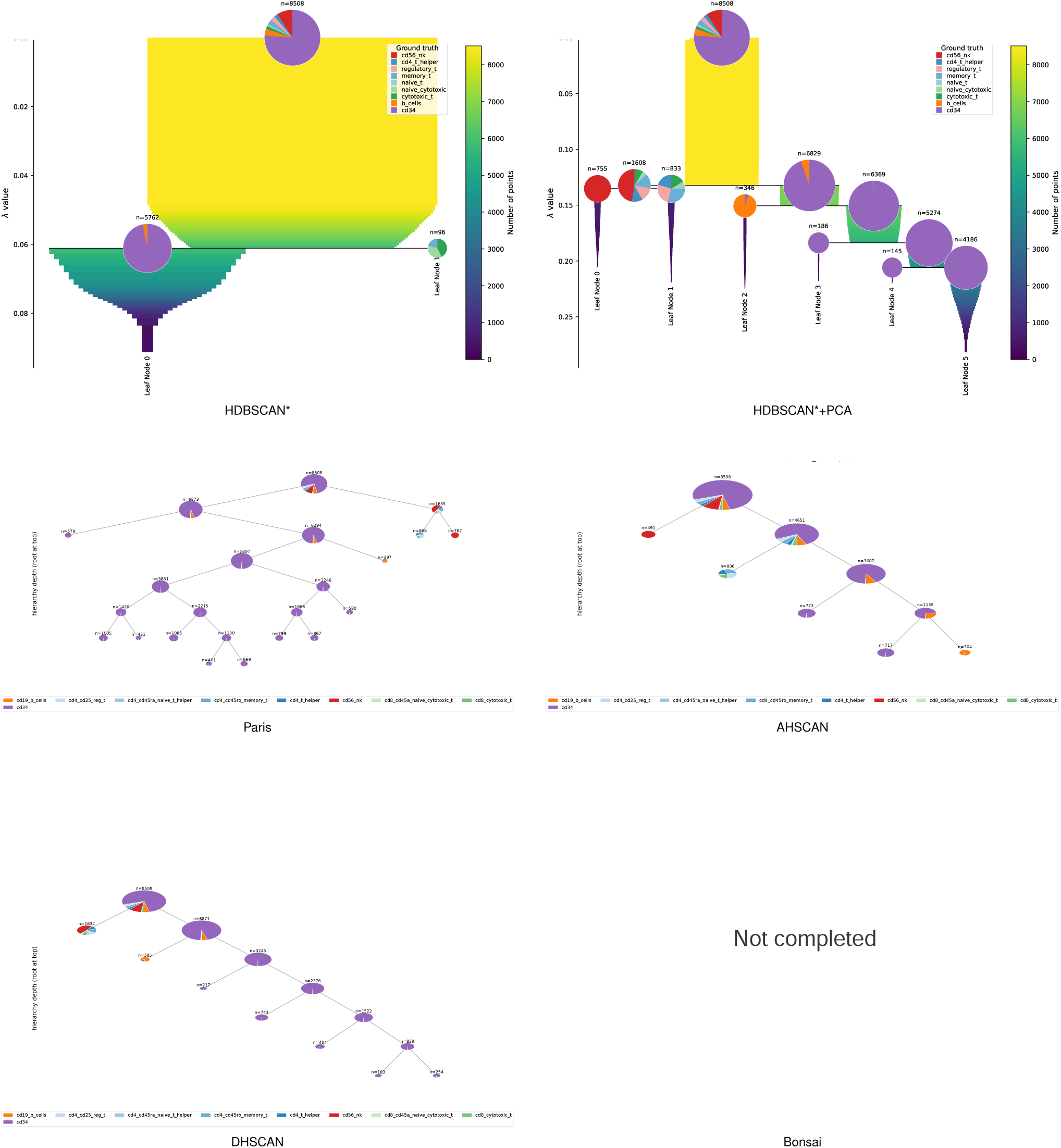
Hierarchical clustering results for the Zheng dataset obtained with HDBSCAN*, HDBSCAN*+PCA, Paris, AHSCAN, and DHSCAN. No Bonsai hierarchy is shown because the available implementation did not complete within one week of runtime. HDBSCAN* produces an overly coarse hierarchy. HDBSCAN*+PCA introduces additional subdivisions, allowing it to detect CD56 NK cells (red), B cells (orange) and CD34+ cells (purple), but T-cell populations (blue/green) remain mixed. Paris and AHSCAN are able to detect clusters related to NK cells (red), B cells (orange) and T cells (blue/green). DHSCAN produces similar results, but did not differentiate between NK and T cells.

**Figure S6.**
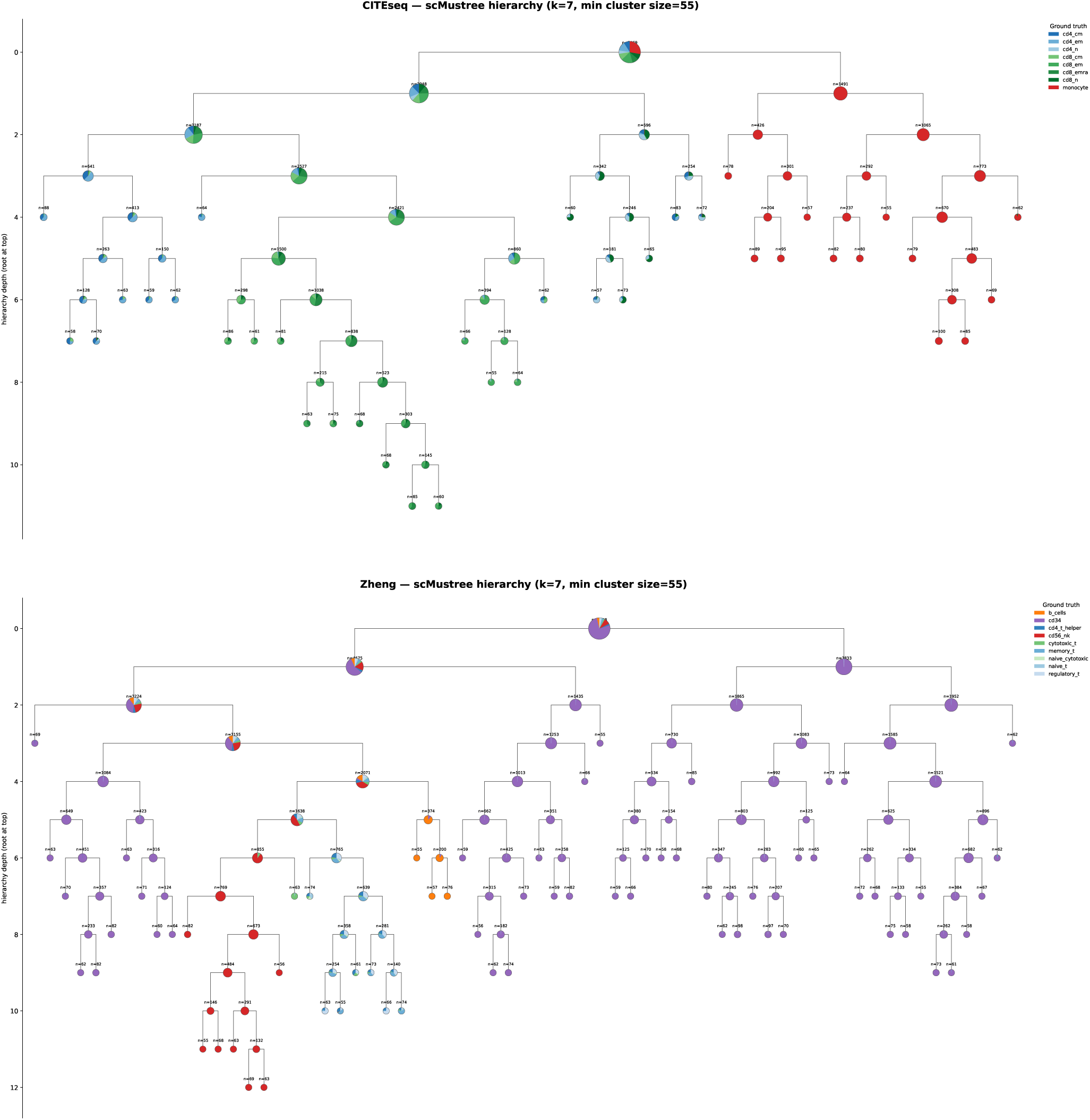
Hierarchical clustering results obtained with scMustree for the CITE-seq and Zheng datasets. For CITE-seq, scMustree clearly separates the monocyte-dominated branch (red) from the T-cell populations, although several CD4 (blue)/CD8 (green) subpopulations remain mixed at the middle branch. For Zheng, the hierarchy is considerably more detailed, but one major population (CD34; purple) dominates many internal branches and appears in several major branches on both sides of the tree.

**Figure S7.**
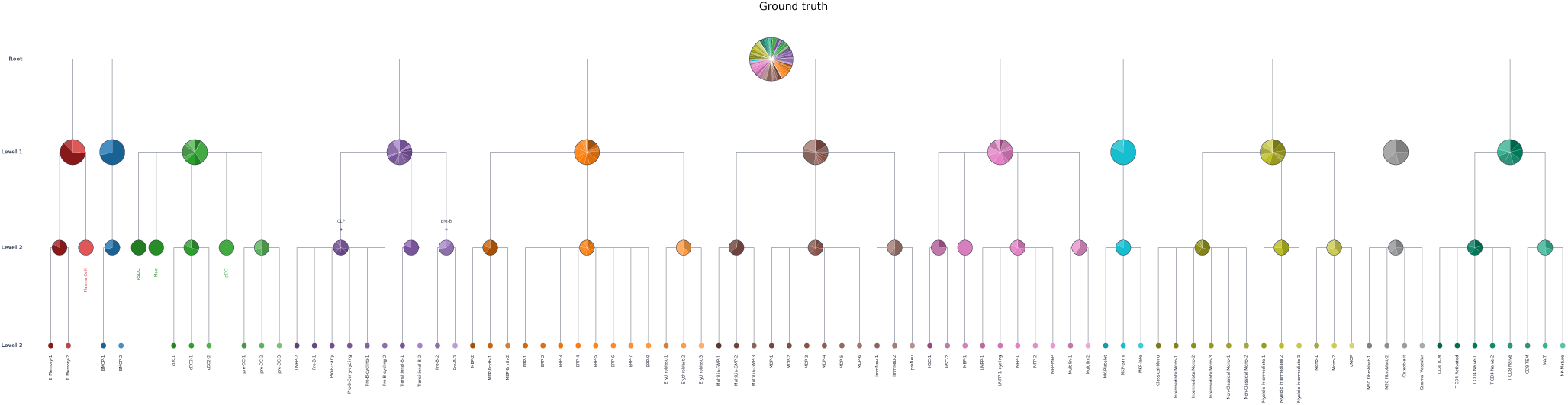
Biological reference hierarchy for the Bone-marrow dataset. The annotation contains three levels of organization spanning multiple mature and progenitor cell populations. Colors indicate the biological reference populations and are used consistently in the corresponding inferred-hierarchy visualizations.

**Figure S8.**
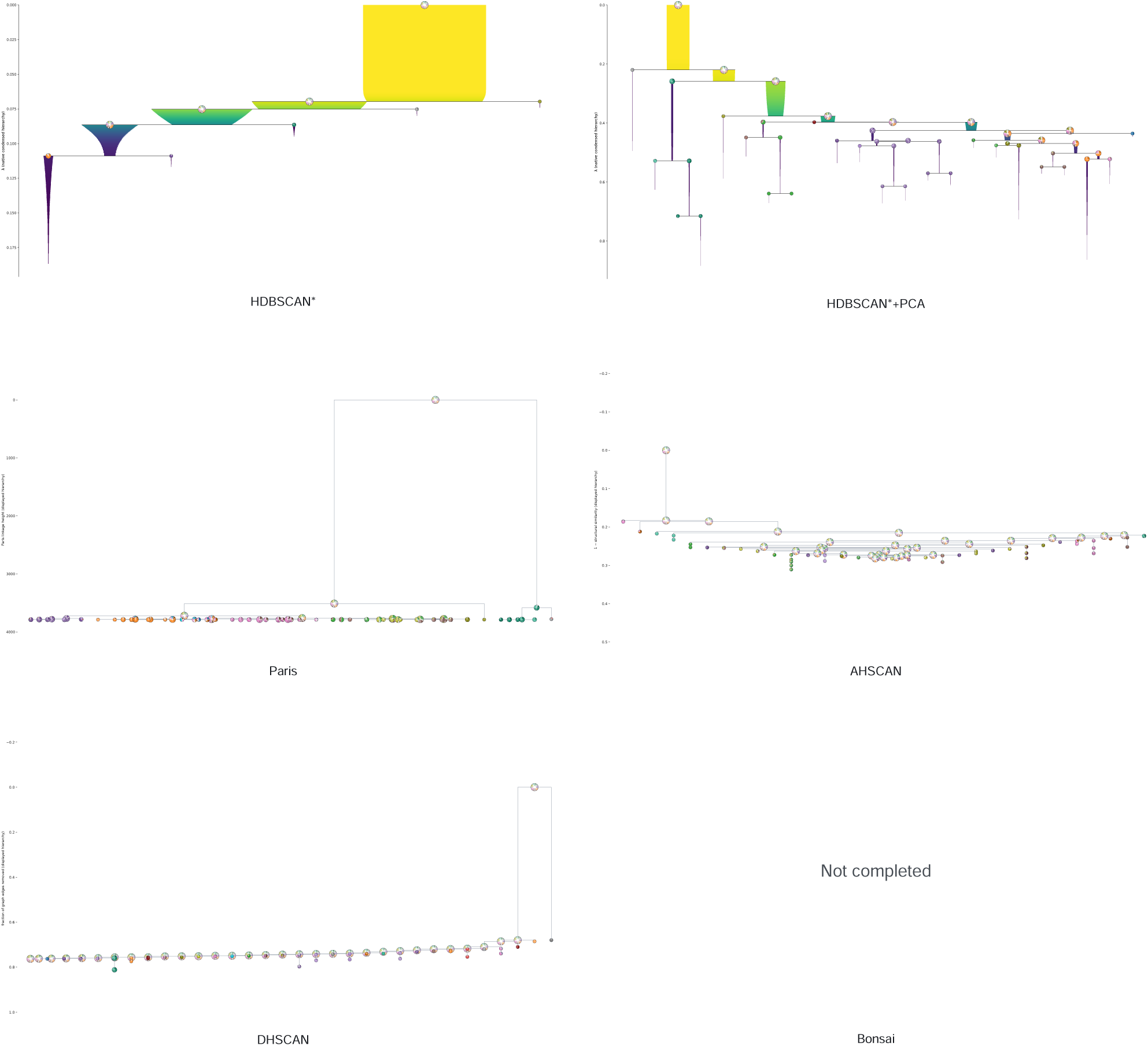
Hierarchical clustering results for the Bone-marrow dataset obtained with HDBSCAN*, HDBSCAN*+PCA, Paris, AHSCAN, and DHSCAN. See Supplementary Fig. S7 for color legend. HDBSCAN* produces a comparatively coarse hierarchy that captures little of the fine-grained reference organization. PCA generates substantially more internal structure, but considerable mixing remains across branches. Paris produces a largely shallow organization with most populations separated only near the leaves, while AHSCAN and DHSCAN yield predominantly chain-like or weakly nested structures that do not clearly reproduce the multi-level biological organization. No Bonsai hierarchy is shown because the available implementation did not complete within one week of runtime.

**Figure S9.**
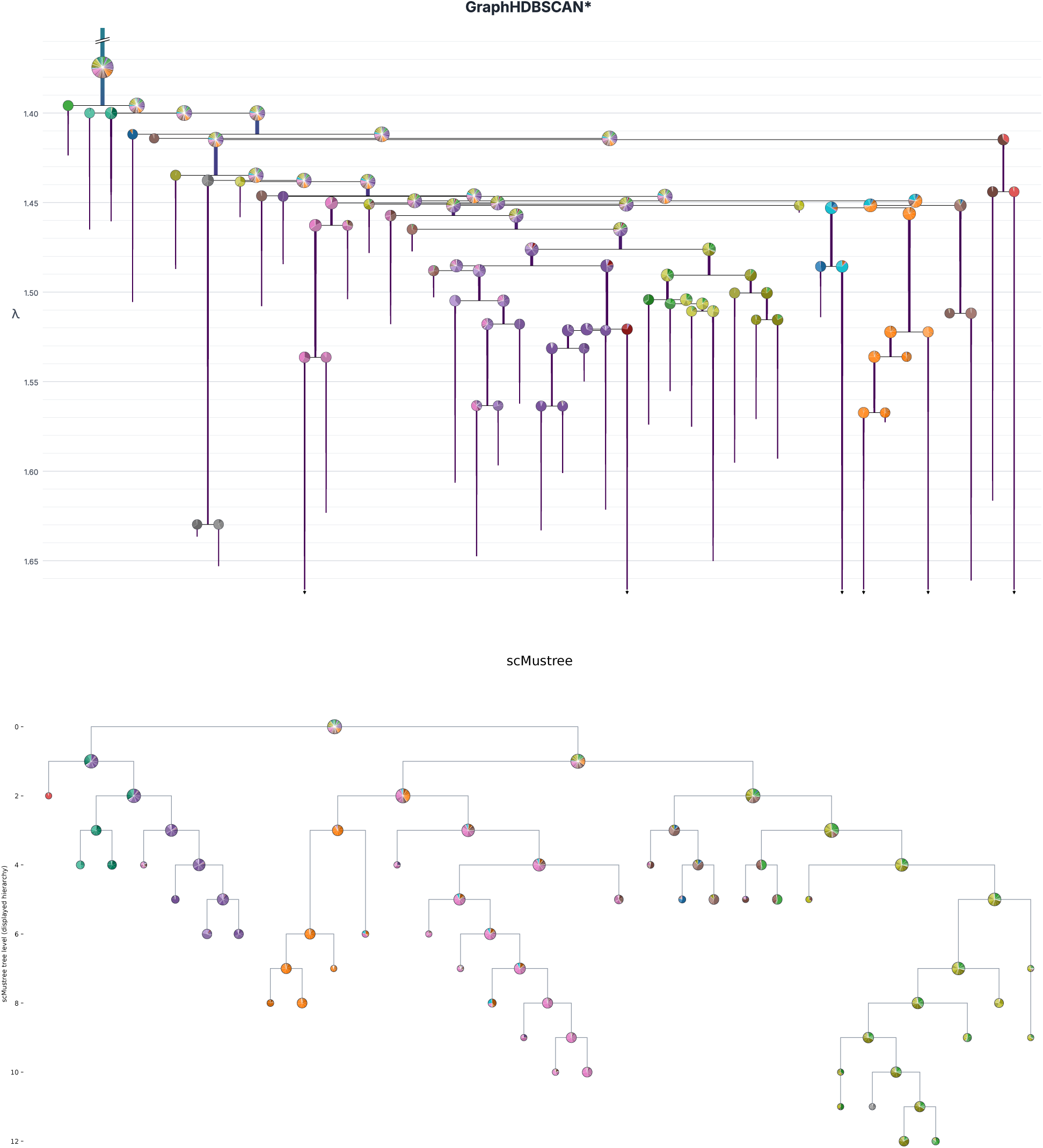
Hierarchical clustering results for GraphHDBSCAN* and scMustree on the Bone-marrow dataset. See Supplementary Fig. S7 for color legend. GraphHDBSCAN* shows extensive multi-level organization, with branches progressively becoming enriched for individual reference populations as the hierarchy is refined. scMustree also produces a detailed hierarchy, but several internal branches remain mixed (right main branch) and individual reference populations are distributed across multiple parts of the tree. For visualization, a minimum displayed cluster size of 55 cells was used to suppress very small branches and improve readability.

