## Supplementary_Data_sc2marker_Complete_results for "GraphHDBSCAN*: Graph-based Hierarchical Clustering on High-dimensional Spaces with Applications to Single-cell RNA Sequencing Data": Supplementary_Data_1_sc2marker_CITEseq.html


### sc2marker

###### 22 January, 2026

### Cluster: -1

#### Table

```
celltype <- "-1"
markers <- markers.all[[celltype]]
get_antibody(markers, org = "human")
```

#### Ridge

```
plot_ridge(scrna, id = celltype, genes = markers[1:6,]$gene, 
           aggr.other = FALSE, ncol = 3, assay = "RNA", slot = "data")
```

### Cluster: 3

#### Table

```
celltype <- "3"
markers <- markers.all[[celltype]]
get_antibody(markers, org = "human")
```

#### Ridge

```
plot_ridge(scrna, id = celltype, genes = markers[1:6,]$gene, 
           aggr.other = FALSE, ncol = 3, assay = "RNA", slot = "data")
```

### Cluster: 1

#### Table

```
celltype <- "1"
markers <- markers.all[[celltype]]
get_antibody(markers, org = "human")
```

#### Ridge

```
plot_ridge(scrna, id = celltype, genes = markers[1:6,]$gene, 
           aggr.other = FALSE, ncol = 3, assay = "RNA", slot = "data")
```

### Cluster: 0

#### Table

```
celltype <- "0"
markers <- markers.all[[celltype]]
get_antibody(markers, org = "human")
```

#### Ridge

```
plot_ridge(scrna, id = celltype, genes = markers[1:6,]$gene, 
           aggr.other = FALSE, ncol = 3, assay = "RNA", slot = "data")
```

### Cluster: 2

#### Table

```
celltype <- "2"
markers <- markers.all[[celltype]]
get_antibody(markers, org = "human")
```

#### Ridge

```
plot_ridge(scrna, id = celltype, genes = markers[1:6,]$gene, 
           aggr.other = FALSE, ncol = 3, assay = "RNA", slot = "data")
```

### Cluster: 4

#### Table

```
celltype <- "4"
markers <- markers.all[[celltype]]
get_antibody(markers, org = "human")
```

#### Ridge

```
plot_ridge(scrna, id = celltype, genes = markers[1:6,]$gene, 
           aggr.other = FALSE, ncol = 3, assay = "RNA", slot = "data")
```
