## Supplementary_Data_sc2marker_Complete_results for "GraphHDBSCAN*: Graph-based Hierarchical Clustering on High-dimensional Spaces with Applications to Single-cell RNA Sequencing Data": Supplementary_Data_2_sc2marker_Zheng.html


### sc2marker

###### 23 January, 2026

### Cluster: cluster 5

#### Table

```
celltype <- "cluster 5"
markers <- markers.all[[celltype]]
get_antibody(markers, org = "human")
```

#### Ridge

```
plot_ridge(scrna, id = celltype, genes = markers[1:6,]$gene, 
           aggr.other = FALSE, ncol = 3, assay = "RNA", slot = "data")
```

### Cluster: cluster -1

#### Table

```
celltype <- "cluster -1"
markers <- markers.all[[celltype]]
get_antibody(markers, org = "human")
```

#### Ridge

```
plot_ridge(scrna, id = celltype, genes = markers[1:6,]$gene, 
           aggr.other = FALSE, ncol = 3, assay = "RNA", slot = "data")
```

### Cluster: cluster 4

#### Table

```
celltype <- "cluster 4"
markers <- markers.all[[celltype]]
get_antibody(markers, org = "human")
```

#### Ridge

```
plot_ridge(scrna, id = celltype, genes = markers[1:6,]$gene, 
           aggr.other = FALSE, ncol = 3, assay = "RNA", slot = "data")
```

### Cluster: cluster 1

#### Table

```
celltype <- "cluster 1"
markers <- markers.all[[celltype]]
get_antibody(markers, org = "human")
```

#### Ridge

```
plot_ridge(scrna, id = celltype, genes = markers[1:6,]$gene, 
           aggr.other = FALSE, ncol = 3, assay = "RNA", slot = "data")
```

### Cluster: cluster 3

#### Table

```
celltype <- "cluster 3"
markers <- markers.all[[celltype]]
get_antibody(markers, org = "human")
```

#### Ridge

```
plot_ridge(scrna, id = celltype, genes = markers[1:6,]$gene, 
           aggr.other = FALSE, ncol = 3, assay = "RNA", slot = "data")
```

### Cluster: cluster 2

#### Table

```
celltype <- "cluster 2"
markers <- markers.all[[celltype]]
get_antibody(markers, org = "human")
```

#### Ridge

```
plot_ridge(scrna, id = celltype, genes = markers[1:6,]$gene, 
           aggr.other = FALSE, ncol = 3, assay = "RNA", slot = "data")
```

### Cluster: cluster 0

#### Table

```
celltype <- "cluster 0"
markers <- markers.all[[celltype]]
get_antibody(markers, org = "human")
```

#### Ridge

```
plot_ridge(scrna, id = celltype, genes = markers[1:6,]$gene, 
           aggr.other = FALSE, ncol = 3, assay = "RNA", slot = "data")
```
